# Multi-region coordination of evidence accumulation and decision commitment in the brain

**DOI:** 10.1101/2024.08.21.609044

**Authors:** Adrian G. Bondy, Julie A. Charlton, Thomas Zhihao Luo, Charles D. Kopec, Wynne M. Stagnaro, Sarah Jo C. Venditto, Kaiyue Shi, Eva Yi Xie, Maxime Beau, Laura Lynch, Sanjeev Janarthanan, Stefan N. Oline, Timothy D. Harris, Carlos D. Brody

## Abstract

Decision-making is thought to involve two phases: first, evidence accumulation, then, decision commitment. Accumulation is represented broadly in the brain, but whether this best matches single versus multiple accumulator models is unknown. Here, analysis of simultaneous, bilateral recordings across sensory, association, and motor regions, in both cortex and subcortical structures, strongly supports a single accumulator, and suggests that accumulated noise (diffusion) originates frontally. Decision commitment can occur internally, without an overt report. Long-standing competing models of covert commitment disagree as to whether it is abrupt or continuous. Temporally aligning data on a single-trial estimator of internal, covert commitment (“nTc”) revealed a rapid (≤50 ms) drop in the accumulator’s sensitivity to incoming sensory evidence, favoring abrupt transition models, and indicating a discrete cross-brain state change at nTc that was first detectable in motor regions. These data discriminate between decades-old models of accumulation, and between decades-old models of the transition to commitment.

## Introduction

Decision making is the cognitive process of selecting one out of a set of options, based on available evidence. Decisions are commonly conceived as divided into two sequential phases: first, evidence accumulation (also thought of as deliberation); second, decision commitment, which corresponds to the subject making up their mind and no longer considering new evidence. Notably, evidence accumulation and commitment are both concepts from mathematical psychology. How the two map onto the brain remains unclear: the functional circuit architecture underlying each phase, and the neural dynamics that demarcate the transition between them, remain largely unresolved.

Multiple studies, in several species, have consistently shown that similar representations of the subject’s evolving decision can be found in surprisingly widespread regions of the brain, including cortex^1–11^, basal ganglia^12,13^, midbrain^14,15^, and brainstem^16,17^ The most immediate interpretation is that this reflects a single, integrated decision formation process^17–19^. However, similar representations may arise from separate, entirely independent, functional circuits (see **Figure S1** for network models constructed to demonstrate this point). Indeed, some long-standing decision making models rely on multiple separate evidence accumulators, with choices arising from a combination of accumulators for distinct options^20–27^, using distinct accumulation algorithms^28^, or using distinct evidence sources^5,29^. Whether neural activity matches a single accumulator better, or one of these multiple accumulator variants better, and how such computations map onto functional circuits in the brain, is not yet known (**Figure 1a**).

**Figure 1.**
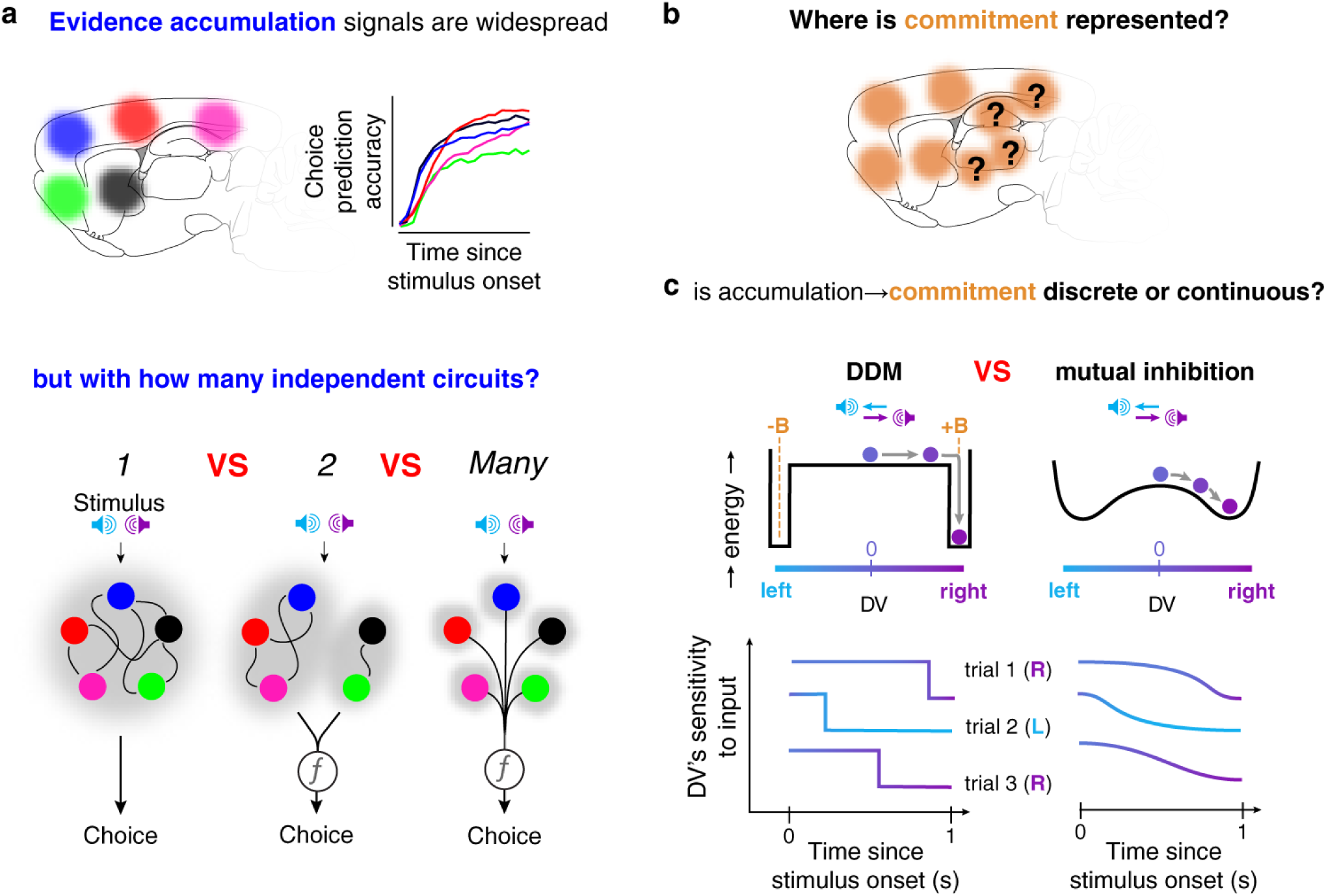
Unresolved questions about the neural representation of evidence accumulation and covert decision commitment. **a.** (Top) Many studies show that averaged over trials, numerous regions across the brain show a gradually increasing ability to predict a subject’s upcoming. But this is consistent with longstanding single-as well as multiple-accumulator models of decision-making. (Bottom) Brain regions could form functional circuits embodying one, two, or many accumulators. Colored circles indicate individual regions; larger gray ellipses demarcate independent accumulators. The function *f* represents an unknown mechanism for combining decision-related activity from different regions to generate a behavioral choice. **b.** Whether or not covert commitment is represented broadly in the brain, and where it originates, is unknown. **c.** Long-standing competing classes of models propose that the transition from accumulation to commitment is either an abrupt change of state (left, DDM and its variants) or a graded, continuous change from one end to the other of a single spectrum (right, nonlinear mutual inhibition). (Top) “Energy” landscape for the two models, with the evolution of the decision variable on an example trial shown. The system tends to evolve downhill in the energy landscape. Right(Left) clicks push the system rightwards(leftwards). Noise in the dynamics is assumed present but is not shown in these illustrative cartoons. (Bottom left) The DDM predicts that the sensitivity of the decision variable DV to incoming sensory input will drop abruptly when the subject covertly commits, an event that happens at different times on different trials when the DV reaches one of the absorbing bounds (+B or −B). (Bottom right). Nonlinear mutual inhibition predicts a gradual change in sensitivity on individual trials. When averaging across trials without first aligning time to the moment of commitment, both models predict a gradual drop in the trial-averaged sensitivity.

The logic that nodes in a connected circuit will tend to co-fluctuate with each other has led to using patterns of neural activity co-fluctuations to delineate functional circuits^30–34^. Neural “decision variables”, DV(t)s^2,10,35–38^, the linear weighting of the activity of tens to hundreds of simultaneously-recorded neurons that best predicts the animal’s upcoming choice, were introduced to provide moment-to-moment, single-trial temporal resolution in the estimated state of the evidence accumulation process and its fluctuations^10,35,39^. Nevertheless, simultaneous measurement of neural DV(t)s across many regions during gradual evidence accumulation, and the ensuing functional circuit analysis of their co-fluctuations, has not yet been attempted. We therefore set out to simultaneously obtain neural DV(t)s across sensory, association, and motor regions, across both hemispheres, and across both cortical and subcortical regions, by developing methods to chronically implant eight Neuropixels 1.0 probes^40^ in freely moving rats. Using these, we simultaneously recorded up to 4,032 neurons across up to 20 regions per session while the rats performed a well-characterized sensory evidence accumulation task, the “Poisson Clicks” task^6,41,42^. We found that a functional circuit analysis of these data provides strong evidence favoring single, not multiple accumulator models.

Subsequent to evidence accumulation, the transition to decision commitment can occur in two ways. *Overt* commitment directly leads to action: a baseball player’s decision about whether or not to swing at a pitch is immediately expressed in their actions. In contrast, in *covert* commitment, the subject settles on a choice but only expresses it as an action later: e.g., halfway through a boring undergraduate lecture, you decide that, after you get home, you’ll drop the class. By temporally disentangling commitment from motor initiation, tasks that elicit covert commitment provide a critical advantage: the time period after covert commitment but before the overt decision report allows studying how commitment *per se* affects neural processing, without neural activity being curtailed by the end of the experimental trial or overtaken by an overt motor response. Nevertheless, covert commitment is difficult to study, because by its very nature, its timing is hidden. Consequently, little is known about where in the brain covert commitment is represented (**Figure 1b**) and about the neural dynamics that underlie it. A class of prominent mathematical psychology models, including the drift diffusion model^43^ (DDM), proposes that covert commitment corresponds to a discrete state change, triggered when the amount of accumulated evidence reaches a bound. This contrasts with a class of also prominent and longstanding competing models, nonlinear mutual inhibition models^44–47^, which propose instead that accumulation and covert commitment are two ends of a graded continuum (**Figure 1c**), thus raising the fundamental question as to whether accumulation and commitment are truly distinct phases. Lacking an estimate of the moment of covert commitment on each trial, most existing studies of the relationship between cellular-resolution neural activity and covert commitment^10,48,49^ have provided only indirect evidence for commitment, and are equally consistent with abrupt versus continuous classes of models.

A recent study^50^, using large-scale recordings in rats performing the same task we use here, developed a method to use neural activity in a set of frontal regions to estimate whether covert commitment occurred, and most importantly, its timing on each trial (labeled “nTc” for “neurally-inferred time of commitment”). Combined with our large-scale recordings, nTc presents a powerful new opportunity to probe the cross-brain functional circuits underlying covert commitment, and assess whether the neural representation of accumulated evidence changes at covert decision commitment in a way that is abrupt or graded. We applied the nTc estimation method to data from each recorded region, thus obtaining simultaneous single-trial estimates of DV(t)s and nTc across a large set of brain regions. Analyses of these data strongly favor the abrupt transition class of models, and show that while covert commitment is broadly represented across the brain, it appears to originate in motor regions.

### Multi-region simultaneous electrophysiology during perceptual decision-making

We developed new methods for large-scale, chronic electrophysiological recording using 8 Neuropixels 1.0 probes per rat^40^. We chose 4 penetration sites, each mirrored bilaterally, for a total of 8 probe insertions per subject. Successful implantation of probes in this dense configuration required the design of novel implantation hardware, an accurate CAD model of the entire assembly in relation to the skull, and new protocols for the lengthy and invasive surgery (see **Methods** and **Figure S3**). We used brain clearing and lightsheet imaging to visualize the implanted probe tracks (**Figure 2b,c**) and combined this with electrophysiological signatures to confirm probe placement. For visualization, we registered the probe tracks to the Princeton RAtlas^51^ (**Figure S2**).

**Figure 2.**
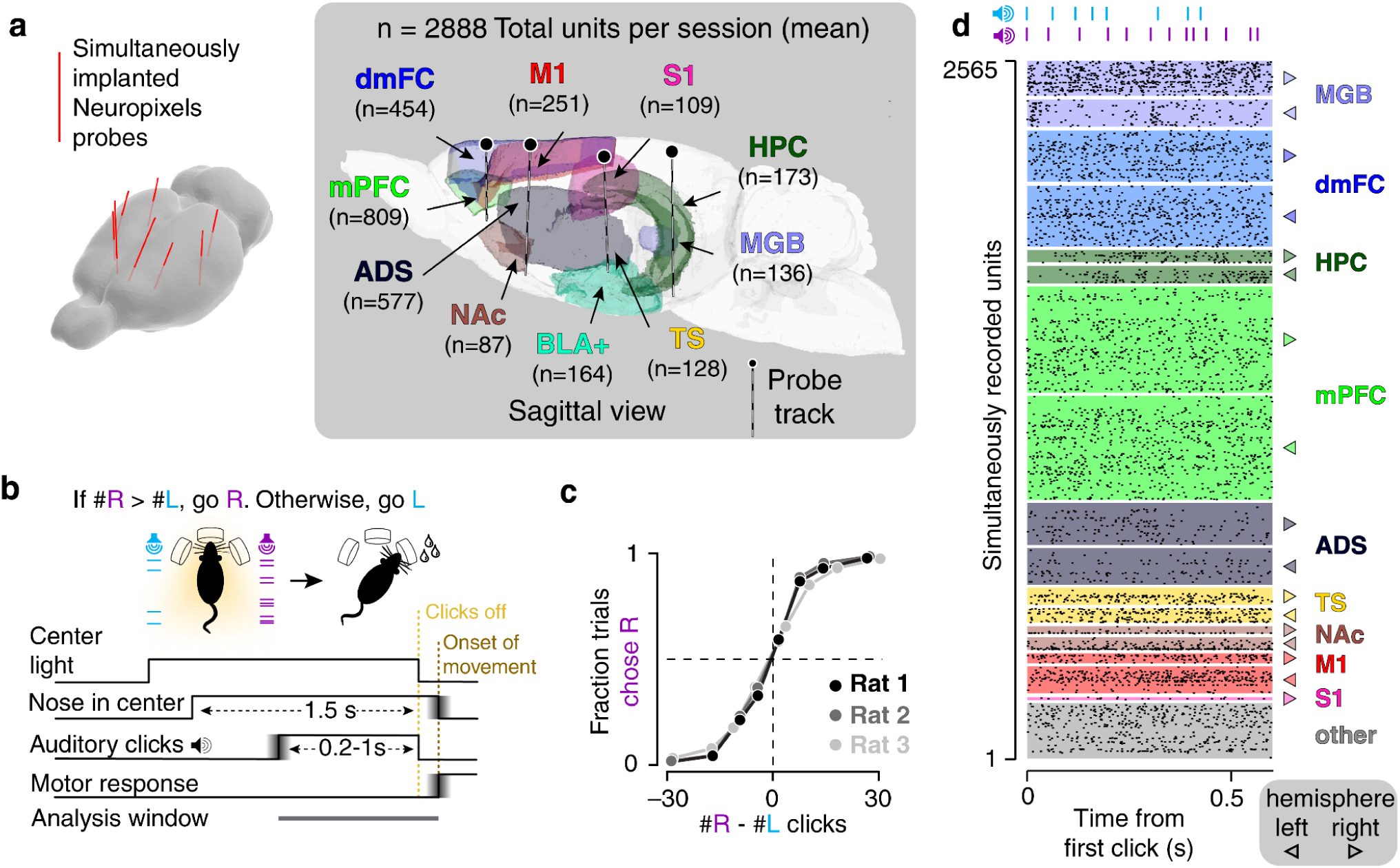
Brain-wide simultaneous electrophysiology during perceptual decision-making. **a.** (left) Schematic showing the arrangement of simultaneously implanted Neuropixels 1.0 probes in a rat brain. (right) Simultaneous placement of 8 chronically implanted Neuropixels probes, with average recorded unit counts per region across sessions (medial prefrontal cortex, mPFC; primary motor cortex, M1; somatosensory cortex, S1; dorsomedial frontal cortex, dmFC; anterior dorsal striatum, ADS; nucleus accumbens, NAc; tail of striatum, TS; basolateral amygdala, BLA+; hippocampus, HPC; medial geniculate body, MGB). **b**. Schematic of a single trial in the “Poisson Clicks” behavioral task. On each trial, at the end of two simultaneous trains of randomly timed auditory clicks, subjects are rewarded for orienting to the side that had the greater total number of clicks. Gradients represent variable timing in the task timelines. **c**. Psychometric functions. **d**. Spike raster of simultaneously recorded neural responses across the brain on a single trial.

Each recording session (3 rats, 19 sessions, 6-7 sessions per rat) yielded thousands of simultaneously recorded units (median=2,888, range 2,385-4,032) from tens of brain regions (median = 19, range 13-20; see **Figure S4** for per-session information). We focused on a subset of the regions recorded, chosen for their known or potential involvement in the perceptual decision-making task described below: dorsomedial frontal cortex (dmFC)^50^, medial prefrontal cortex (mPFC), primary motor cortex (M1), primary somatosensory cortex (S1), anterior dorsal striatum (ADS)^12^, the tail of the striatum (TS)^52^, the hippocampus (HPC), basolateral amygdala (and inclusion of nearby nuclei; BLA+), nucleus accumbens (NAc), and medial geniculate body (MGB). See **Table S1** for further details.

ADS and dmFC (the strongest cortical input source to ADS) have been directly linked, through recording and causal perturbation, to evidence accumulation during the task used here^12,50^. dmFC is bidirectionally connected^53^ to M1 and mPFC, and ADS receives some input from those two regions^50,54^. MGB is the principal relay for all auditory information reaching the forebrain^55^, and projects directly to TS^56,57^, which has been shown to be necessary for auditory decision-making^58,59^. S1 is reciprocally connected to M1^60^, projects broadly within striatum^54^, and has recently been established to play a causal role in the formation of perceptual decisions^61^. Decision-related neurons have also been discovered in HPC and NAc during accumulation-of-evidence tasks^50,62^. BLA+ contains many auditory-responsive neurons^63^ and projects broadly throughout the striatum^54^.

Before the implantation surgery, the subjects were trained to perform an established auditory decision-making task requiring accumulation of evidence presented in the form of randomly-timed pulsatile auditory clicks^41,41,64^ (the “Poisson Clicks” task; **Figure 2b**). Subjects initiate a trial by poking their nose into a center nose port, and keeping it there; after a variable delay (0.5 - 1.3 s), two randomly-timed trains of auditory clicks are played from speakers to the subject’s left and right, until a visual “go” cue, 1.5 s after the initial center poke. The subject is then free to withdraw from the center port, poke its nose into one of two side ports, and receives reward if it poked into the side that had played the greater total number of clicks. The optimal strategy thus requires gradually accumulating clicks over time on each trial. After surgery, all three rats continued to exhibit good behavioral performance (see **Methods**). We focus our analyses on the period from the start of the clicks to the end of the clicks or the withdrawal from the center poke, always before the animal responds.

### A single evidence accumulation process is shared across the brain

We used neural decision variables DV(t), an established method introduced to estimate the moment-to-moment evolution of the decision on single trials^2,10,35–38,65^. We took firing rate data from all simultaneously recorded neurons in a region, time-aligned to the start of the sensory stimulus, and at each point in time, used logistic regression to determine the weighted sum of neural firing rates that best predicted the animal’s choice across all trials (**Figure 3a**). Taken as a vector, the set of weights define a direction in neural space sometimes referred to as the “choice axis” or the “choice coding direction”^65–67^. For each trial and each time point, this weighted sum yields the “decision variable” or DV(t)^35^ (**Figure 3b**; see **Methods**). Since it is compiled across many noisy neurons, DV(t) has higher resolution than signals obtained from individual neurons^35^. A more positive(negative) value of DV(t) corresponds to a higher logistic regression probability of a rightward(leftward) choice.

**Figure 3.**
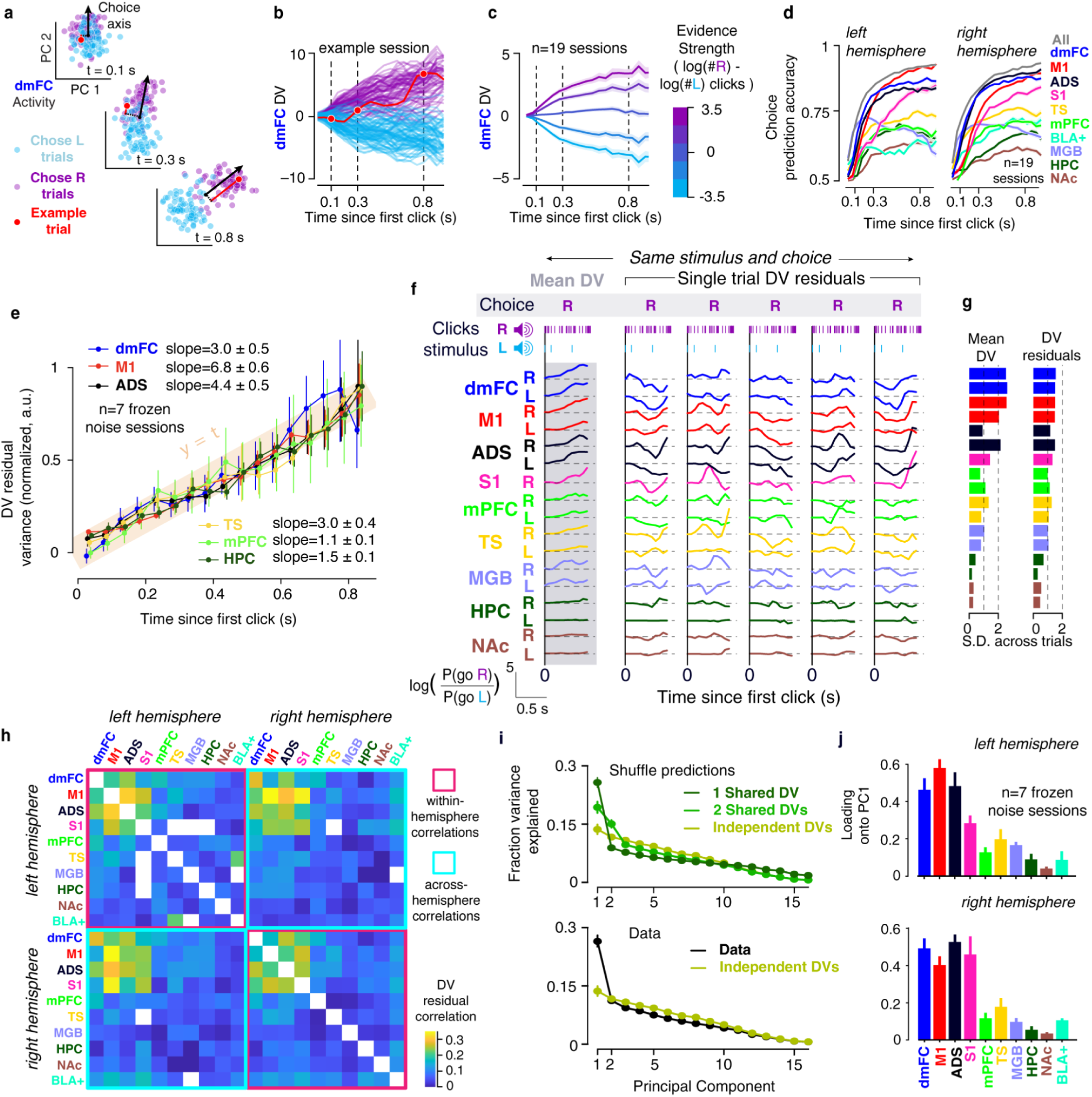
A single evidence accumulation process is shared across the brain. **a.** Single trial dmFC population activity, shown separately at three time points relative to the first click, in a two-dimensional state space, for one example session. Blue (purple) dots depict neural activity on chose L (chose R) trials. Black arrows indicate the weight vectors that best predict the subject’s choice using logistic regression, whose direction defines a “choice axis”. Red lines indicate projection onto the weight vector (the “decision variable”) for an example trial. **b**, Decision variable time course – DV(t) – across all trials; the example trial is highlighted in red. **c,** Time course of trial-averaged DV(t)s, conditioned on given signed trial difficulties. Error bar is +/−1 s.e., hierarchical bootstrap across sessions.**d**, Time course of choice prediction accuracy using neural activity from each brain region and from all regions combined. Mean +/−1 s.e. across sessions shown. **e**. Across-trial variance of DV(t) residuals increases linearly over time. Error bars indicate +/−1 s.e. across frozen noise seeds. Only timepoints up to 200 ms before each trial’s neurally estimated time of commitment (nTc-all; using all recorded neurons) were used (see **Methods**). To focus on comparing linearity across regions, per-region data was shifted and scaled vertically so linear fits to each region all had an intercept of 0 and a slope of 1. Actual slopes for each region are as indicated, in units of log-odds squared per second. Four other regions had data that systematically deviated from a straight line (shown in **Figure S5d**). **f**. Simultaneously obtained DV(t)s for all recorded regions in an example “frozen noise” session, shown for a subset of trials that shared the same click train and choice. The left column shows the average DV(t) across this set of trials for each region and hemisphere, and the remaining columns show the residual fluctuations (DV(t) residuals) for five example trials around that mean. **g**. Standard deviation of the mean and residual (i.e. stimulus- and choice-conditioned mean subtracted) DV(t)s across all trials in the session whose data is shown in b, c and e. **h**, Matrix of DV(t) residual correlations (Pearson’s *ρ*) between pairs of regions, across timepoints and trials within a session. Average across the 7 frozen noise sessions shown. Only timepoints up to each trial’s neurally estimated time of commitment (nTc-all; using all recorded neurons) were used (see Methods). **i**, Scree plot obtained from applying principal components analysis to the set of across-region residual DV(t)s. Each region/hemisphere pair is treated separately. 16 PCs are shown because very few individual sessions had more than 16 region/hemisphere pairs. The top panel displays predictions from the three competing models in Figure 1a, and the bottom panel compares the recorded data with the scenario in which each set of DV(t) residuals are independent, which provides an estimate of the noise floor (see **Methods**). Error bars indicate 1 s.e. obtained from a hierarchical bootstrap across sessions. **j**, Plots showing the loading of each region/hemisphere pair onto the first PC. Error bars indicate 1 s.e. obtained from a hierarchical bootstrap across sessions.

Single-trial DV(t)s in non-human primates have previously been interpreted as reflecting the state of an evidence accumulator^10,35,39^. Several findings, all similar to findings in non-human primates, support that conclusion here with rats. First, DV(t)s on single trials tended to ramp positive on rightward choices and negative on leftward choices^35^ (**Figure 3b**), with the slope of the ramp correlated to the strength of stimulus evidence presented (**Figure 3c**)^10^; Second, DV(t)s quantitatively predicted the fraction of left vs. right choices^10^ (**Figure S5c**); Third, DV(t)s predicted the animal’s choice more accurately than knowledge of the stimulus alone^10^ (**Figure S5b)**. As observed by others^2,16,17,37^, and similarly across left and right hemispheres, we found that the trial-averaged choice prediction accuracy predicted by the DV(t)s was significantly greater than chance, for all recorded regions, and increased gradually over time (**Figure 3d**). The number of neurons recorded per region was only weakly predictive of the region’s choice prediction accuracy (*R*^2^=0.11, p=0.02; **Figure S4b,c**). None of the three regions with the highest choice prediction accuracy (frontal cortical regions M1, dmFC, and anterior dorsal striatum, ADS), all previously implicated in decision-making, had the highest recorded unit count (**Figure S4**), indicating that higher choice coding in these regions was not trivially due to population size.

To control for cross-trial variability due to changes in the stimulus or the subject’s choice, in a subset of 7 sessions, distributed across all 3 rats, we used a “frozen noise” task design. In these frozen noise sessions, only 54 fixed click trains were presented, in random order across trials; each of the 54 trains was thus repeated over multiple trials (median of 10 per session). We then conducted our analyses within sets of trials that had a given choice and click train stimulus, examining residual DV(t) fluctuations relative to the mean DV(t) trajectory within that set. Using DV(t) residuals, two additional findings further supported the conclusion that DV(t)s reflect the state of the evidence accumulator. Decision making models (such as the drift-diffusion model^43^, DDM) posit that the accumulator evolves as a random walk (diffusion), biased by stimulus evidence (drift). When the effect of the stimulus is removed, only diffusion remains, and the accumulator’s variance should increase linearly over time. Linearly increasing variance has therefore been taken as a signature of the accumulator^39,68^. Strikingly, using DV(t) residuals around fixed stimuli, for most recorded regions we found linear growth in across-trial variance up to 0.8 sec after stimulus onset, a period ∼2X longer than previously observed in non-human primates^39,68^ (**Figure 3e**). In the remaining minority of regions, linearity was still a good approximation, despite systematic deviations (**Figure S5d**). DV(t) residuals thus have a strong signature of evidence accumulation. Finally, we ruled out the alternative possibility that single-trial DV(t) residuals are driven by uninstructed spontaneous movements^69,70^. Video analysis showed that uninstructed movements could explain only a small fraction of variance in DV(t) residuals (15% variance explained for M1 DV(t) residuals, and substantially less for other regions; **Figure S6**), and most importantly, DV(t)s from most regions, including the frontal regions with the strongest choice signals (M1, ADS, and dmFC), temporally led fluctuations in uninstructed motor activity (**Figure S6**). This suggests that DV(t)s drive uninstructed movements, and not the other way around, a conclusion consistent with a recent study of mouse secondary motor cortex and uninstructed facial movements^71^.

Having concluded that rat DV(t)s have strong signatures of evidence accumulation, consistent with those found in DV(t)s of monkeys, we turned to examining their cross-region co-fluctuation patterns. The left column of **Figure 3f** shows DV(t)s from 9 brain regions, across both hemispheres, for one example frozen noise session, averaged across repeats of one of the 54 click trains, which is shown at the top. As expected for an easy “go Right” trial, with many more Right clicks than Left clicks, trial-averaged DV(t)s in all brain regions gradually grew increasingly positive as the trial progressed. The five columns to the right show DV(t) residuals from representative single trials. Several striking features, typical of the data as a whole, are immediately apparent: First, the residuals are substantial, with magnitudes comparable to the mean itself (**Figure 3g**). Across regions, the standard deviation of the DV(t) residuals was similar in magnitude to the average DV(t) traces (99% as large as, +/−5% s.e. bootstrap across sessions). Second, the fluctuations are significantly correlated across regions, with different time courses across trials but similar time courses across regions within a trial. Importantly, the co-fluctuations across regions are not due to shared fluctuations in the stimulus driving the DV(t)s or in the subject’s final choice, since these are residual DV(t) correlations: the stimulus is held constant across trials, and all trials used in **Figure 3f** resulted in the same choice (Right). Residual DV(t) correlations between most pairs of regions were well above zero, with the correlations between three regions (dmFC, ADS and M1) clearly standing out from the rest as strongest (*ρ*≈0.3; **Figure 3h),** a finding that was consistent across subjects **(Figure S7**). Notably, this pattern of inter-regional correlations was highly specific to DV(t)s, which project neural activity onto the choice axis: correlations between projections onto other directions, representing other one-dimensional summaries of population activity, such as mean population firing rate, were much lower overall and did not share the same structure (**Figure S8**). Patterns of correlation across left and right hemispheres were strikingly similar to patterns within hemisphere, demonstrating a remarkable degree of inter-hemispheric coordination during evidence accumulation (**Figure 3h**).

Independent accumulators, provided with correlated evidence streams, will produce correlated outputs. But their residuals around a given evidence stream will be independent of each other (**Figure S1**). If different groups of regions reflect different accumulators - for example, if left(right) hemisphere regions were separately accumulating evidence for a right(left) decision, as in race models of accumulation^20–27^ - then we would expect co-fluctuations within but not across groups. More generally, the dimensionality of the set of DV(t) residuals across regions, which we assess here using principal components analysis (PCA), reflects an estimate of the number of independent functional circuits (**Figures S1, 3i**). Examining DV(t) residual correlations across all regions, we found that a single dominant PC accounted for around 27% of the variance in the dataset ([23-30] 95% coverage interval using a hierarchical bootstrap), more than twice the next principal component, which fell below the noise floor (**Figure 3i**; also seen in data from each individual animal, **Figure S7**). While this does not preclude the possibility that a larger dataset would allow further principal components to rise above the noise floor, it is clear that the first component dominated, and DV(t) residual co-fluctuations, across all recorded regions, can be described well by a single scalar variable (the projection onto the first PC). Shuffling analyses showed that the data’s PC spectrum is remarkably similar to what would be expected if all regions reflected a single shared DV (**Figure 3i** and see **Methods**). Dividing each region’s neurons into left-choice-preferring versus right-choice-preferring groups, instead of their hemispheric location, led to the same conclusion (**Figure S9**): left- and right-choice neurons are coupled into a single functional circuit. Overall, these data strongly support single, not multiple, accumulator models of decision-making.

### A frontostriatal subnetwork temporally leads evidence accumulation

To our knowledge, temporal relationships between DV(t)s of different regions have not previously been examined. If signals originate in one set of regions, and are relayed to other regions, we would expect fluctuations in the “receiver” regions to lag those in the “source” regions (**Figure 4a**). Using the frozen noise sessions, and for each pair of regions, we measured their DV(t) residual correlations at different time lags between the regions, resulting in the temporal cross-correlation function of the residuals. Because we are using DV(t) residuals, these correlations exclude those introduced by coding for stimulus or choice. That is, the temporal relationships shown here are separate from and independent of the relative lags with which different regions respond to the stimulus. **Figure 4b** shows the resulting cross-correlation functions for ADS versus all other regions. The functions peak near zero and decay to negligible values within about 0.5 s, reflecting relatively short timescale co-fluctuations, less than the duration of a trial. The peaks are shifted to the left of t=0 for most pairs, indicating that ADS DV(t) residual fluctuations lead those of most other regions.

**Figure 4.**
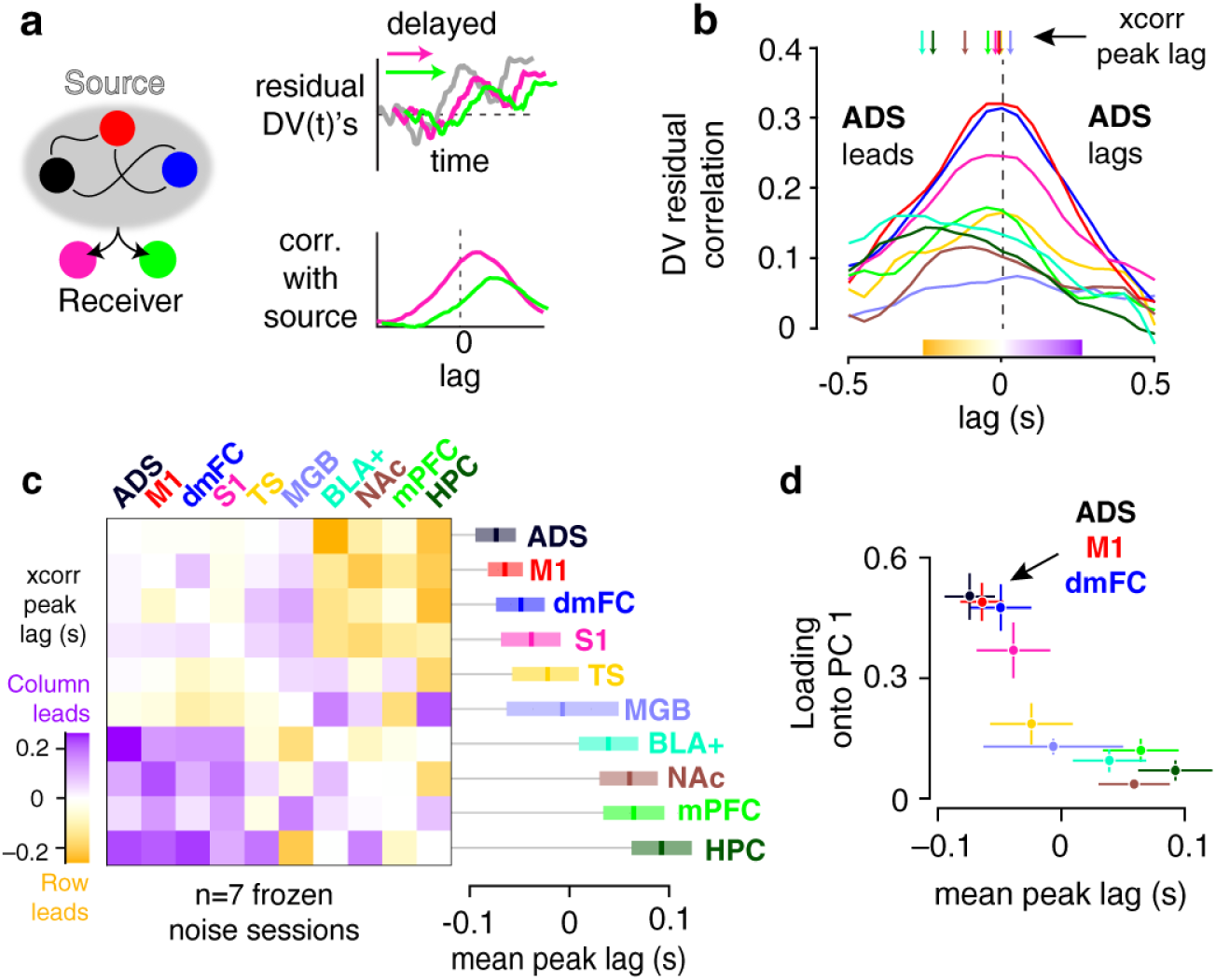
A frontostriatal subnetwork temporally leads evidence accumulation. **a**, Schematic of the motivating problem: identifying where signals related to evidence originate. DV(t)s in source regions are predicted to lead those in recipient regions, leading to shifted cross-correlogram peaks. This should be true even after removing the component related to stimulus and choice (i.e. using DV(t) residuals). **b**, Pearson correlation between the residual DV(t)s of each brain region with ADS as a function of different temporal lags. The lag at which the maximum value occurs (the cross-correlogram peak lag) is shown at the top for each pair. Only timepoints up to each trial’s neurally estimated time of commitment (nTc-all; using all recorded neurons) were used (see Methods). **c**, At left, lag of the cross-correlogram peaks, computed as in panel b, but here shown for all pairs of brain regions. The first row corresponds to the cross-correlogram peaks in (b). Positive values (purple) indicate that the region on the column leads the region on the row. Note that the matrix is therefore antisymmetric by construction. At right, the average of each row in the matrix, with error bars indicating 1 bootstrap s.e. across trials. Negative values indicate a region’s residual DV(t) tends to lead other regions. **d**, Scatter plot of each region’s residual DV(t) loading on PC1 (as in Fig. 2j but mean across hemispheres) versus its mean peak lag in (c).

The left panel of **Figure 4c** shows the results of the peak time lag analysis of **Figure 4b**, but now for all pairs of regions (by rat shown in **Figure S10**). Each region’s peak lag, averaged over all other regions, is shown in the right panel of **Figure 4c**. This shows that during decision formation, ADS, M1 and dmFC – the same set of three interconnected frontostriatal regions that had the strongest choice prediction accuracy in **Figure 3d** – tended to have DV(t) residual fluctuations that led the rest of the brain by the largest amount (∼50 ms). ADS, M1, dmFC were indistinguishable from each other in this measure (using bootstrap test across trials). Using each region’s loading onto the first PC of **Figure 3j** as a measure of the strength with which that region represents the shared accumulation process, and comparing that to the region’s average peak lag, we found that the two were highly correlated (**Figure 4d**). In other words, the strength with which the shared accumulation process is represented is closely correlated with the latency with which it is observed. ADS, M1 and dmFC led in both measures. Taken together, these data suggest that while a single accumulator is being represented across all recorded regions, the accumulator’s value may be computed in a frontal subcircuit, and then broadcast to the rest of the brain.

The fact that DV(t) residuals have a strong signature of the accumulator’s diffusion noise (**Figure 3e**), suggests that their cross-correlations reflect the relative lags of representations of this diffusion noise. To our knowledge, the anatomical source of the accumulator’s diffusion noise (a prevalent component of a wide variety of decision-making models) has not been previously addressed. The results of **Figure 4** specifically suggest frontal regions as the anatomical source of the accumulator’s diffusion noise. (We note that we refer here to the temporally-integrated noise. Our data do not identify the source of the instantaneous noise that, when integrated, becomes the diffusion noise.)

### Cross-brain estimates of the time of covert decision commitment

Covert decision commitment, by definition, is an internal cognitive event, not timelocked to external events such as a stimulus or an overt motor action. Recent work developed a method to use the neural activity of tens or more simultaneously recorded neurons to define the “neurally-inferred time of commitment” or “nTc”^50^ – an estimate of whether and when covert decision commitment occurred on each individual trial.

In brief, the method relies on a model of spiking activity (the “multi-mode drift diffusion model”, MMDDM)^50^ that maps the DDM’s scalar evidence accumulator (labeled here *z*(*t*)) onto high-dimensional firing rates **y**(t) (**Figure 5a**). Each element of **y**(t) represents a recorded neuron’s firing rate, and the accumulator *z*(*t*) is driven by incoming sensory evidence. Critically, the *z*(*t*) → **y**(t) map is allowed to be different during accumulation versus commitment (i.e., before versus after *z*(*t*) hits the absorbing bound); the two maps correspond to different directions in neural space.The abrupt change in neural space direction upon commitment serves as a temporally precise signature of nTc. The model’s parameters are fit to maximize the likelihood of observing the experimental data. Subsequently, for each trial, we compute the posterior distribution of *z*(*t*) given the auditory click times and the recorded spike trains (leaving out the animal’s choice). If this *z*(*t*) posterior distribution reaches a bound with 0.95 probability or greater, the trial is denoted as having covert commitment, and the earliest moment of reaching 0.95 is denoted as the nTc time (**Figure 5b**). Spiking data constrain nTc much more sharply than the sensory stimulus alone (**Figures 5c, S11e**). Two of the key validations of nTc are first, assessing whether or not MMDDM fits held out data better than a single-mode DDM; and second, testing whether the neurally-derived nTc estimates predict the moment in which subject’s choices covertly cease to be affected by incoming sensory evidence. This last is the most critical validation, for it tests an out-of-domain prediction (behavioral choices are not used to determine nTc, but are used in the validation test), and it directly links nTc to the core definition of commitment^50^. We found that the timing of nTc is unlikely to be explained by uninstructed movements: a convolutional neural net trained to distinguish video frames before versus after nTc, did not perform above chance for frames +/−100 ms from nTc (**Figure S12**). We note that here we do not treat MMDDM as a generative model underlying the DV(t)s, which may emerge from more complex dynamics, but only as a tool for inferring commitment timing.

**Figure 5.**
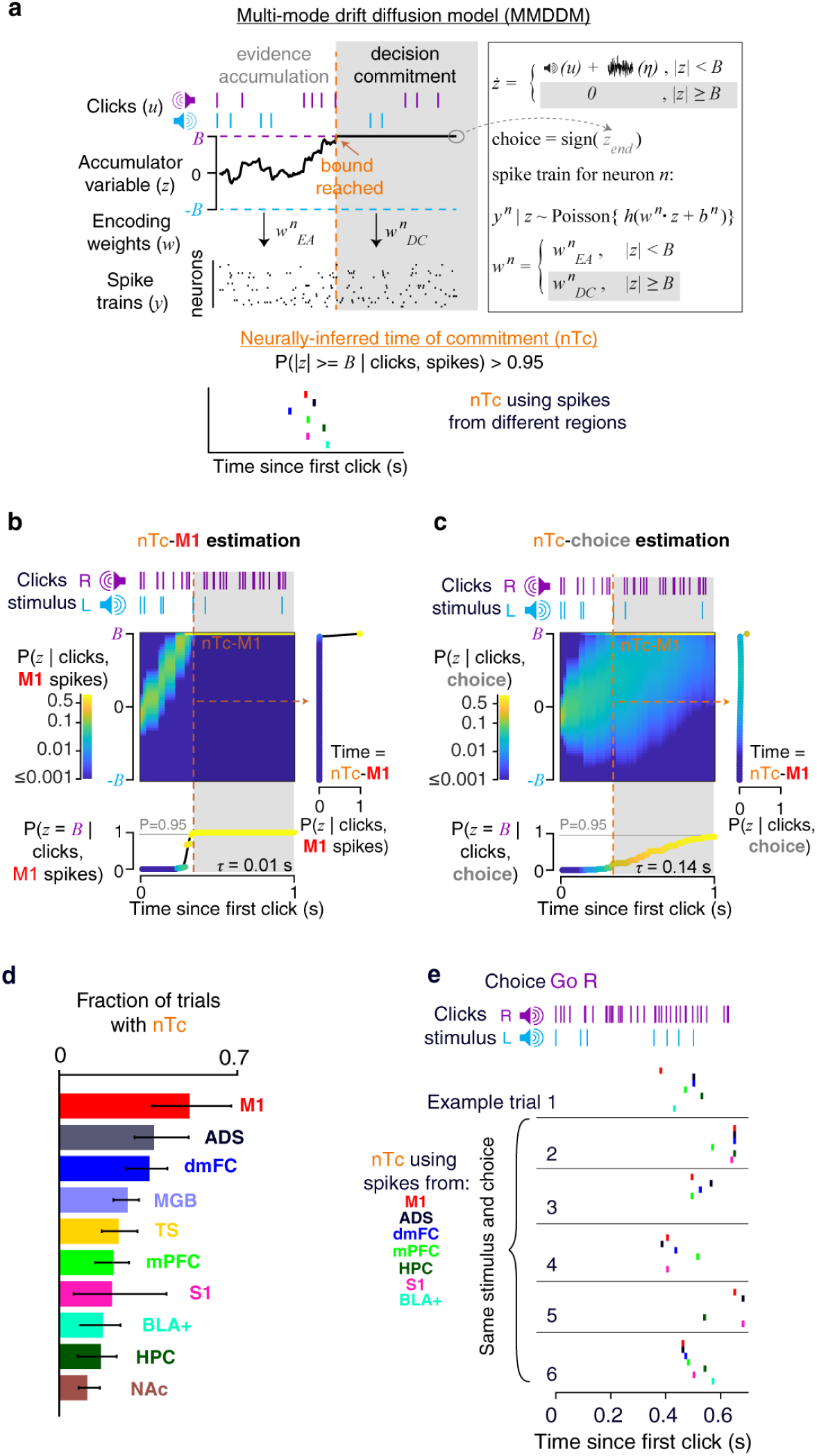
Brain-wide estimates of the time of covert decision commitment. **a**, Schematic illustrating the procedure used to infer region-specific neurally inferred time of commitment (nTc) based on a model assuming accumulation to a decision bound with an abrupt change in neural encoding at commitment. **b**, Posterior probability of commitment over time for an example trial, computed using M1 spike data. **c,** Same trial and format as b, with posterior probability computed using choice. **d**, Detection rates of nTc across brain regions. Error bars indicate hierarchically bootstrapped 95% CI. **e**, Colored dots indicate nTc’s estimated from different brain regions on six example trials with identical click trains and choices, highlighting correlated variability in commitment timing across regions.

The study that developed the nTc estimation method^50^ applied it to neural activity from frontal regions. But posterior and sensory regions were not examined. Applying the nTc method to data from each recorded region here, we found that nTc could be detected most frequently in frontal regions M1 (48% of trials), followed by ADS (35%) and dmFC (33%; **Figure 5d**). Thus, decision commitment signals are strongest in the same set of brain regions that most robustly represent the shared process of decision formation (**Figure 3**). But surprisingly, nTc could be estimated, and fully validated (**Figures S10, S11**), in every recorded region (except NAc, which did not pass the behavioral validation). This included sensory regions like the medial geniculate body (MGB). Covert commitment is thus represented broadly in the brain. Examining frozen noise trials, we found that nTc timing, relative to stimulus onset or offset, was highly variable across trials with identical stimuli and choices (**Figure 5e**). Nevertheless, visual inspection of these frozen noise data immediately suggested that nTc timing is correlated across regions.

### Decision commitment is shared across the brain and is led by M1

Quantifying single-trial temporal relationships between the nTc estimates across regions allows assessing whether they jointly reflect a single moment of commitment for the whole organism as well as where in the brain the commitment signal may first arise. To do this, we followed a procedure similar to our analysis of co-fluctuations amongst DV(t) residuals. We controlled for correlations in nTc timing across regions due trivially to cross-trial changes in stimulus and choice by using frozen noise sessions. For each brain region and for each set of trials with a given click train and choice, we subtracted the mean nTc in those trials, and examined the resulting nTc residuals. To assess the number of independent processes involved in generating nTcs across regions (**Figure 6a**) we used principal components analysis to assess the dimensionality of the space of simultaneously estimated nTc residuals (**Figure 6b**). We tested the approach with synthetic data by running two MMDDM models in parallel, producing simulated spike trains for two disjoint sets of brain regions. Both models were run with identical parameters and were driven by identical stimulus trains. The only difference between them was that each had its own instantiation of noise in its accumulator *z*(*t*). Confirming the analysis method, the two models, conditionally independent given parameters and sensory stimuli, produced two PCs above the noise floor. In contrast, a single MMDDM model for all brain regions produced only one PC above the noise floor (**Figure 6c**, by rat shown in **Figure S13**; see **Methods**).

**Figure 6.**
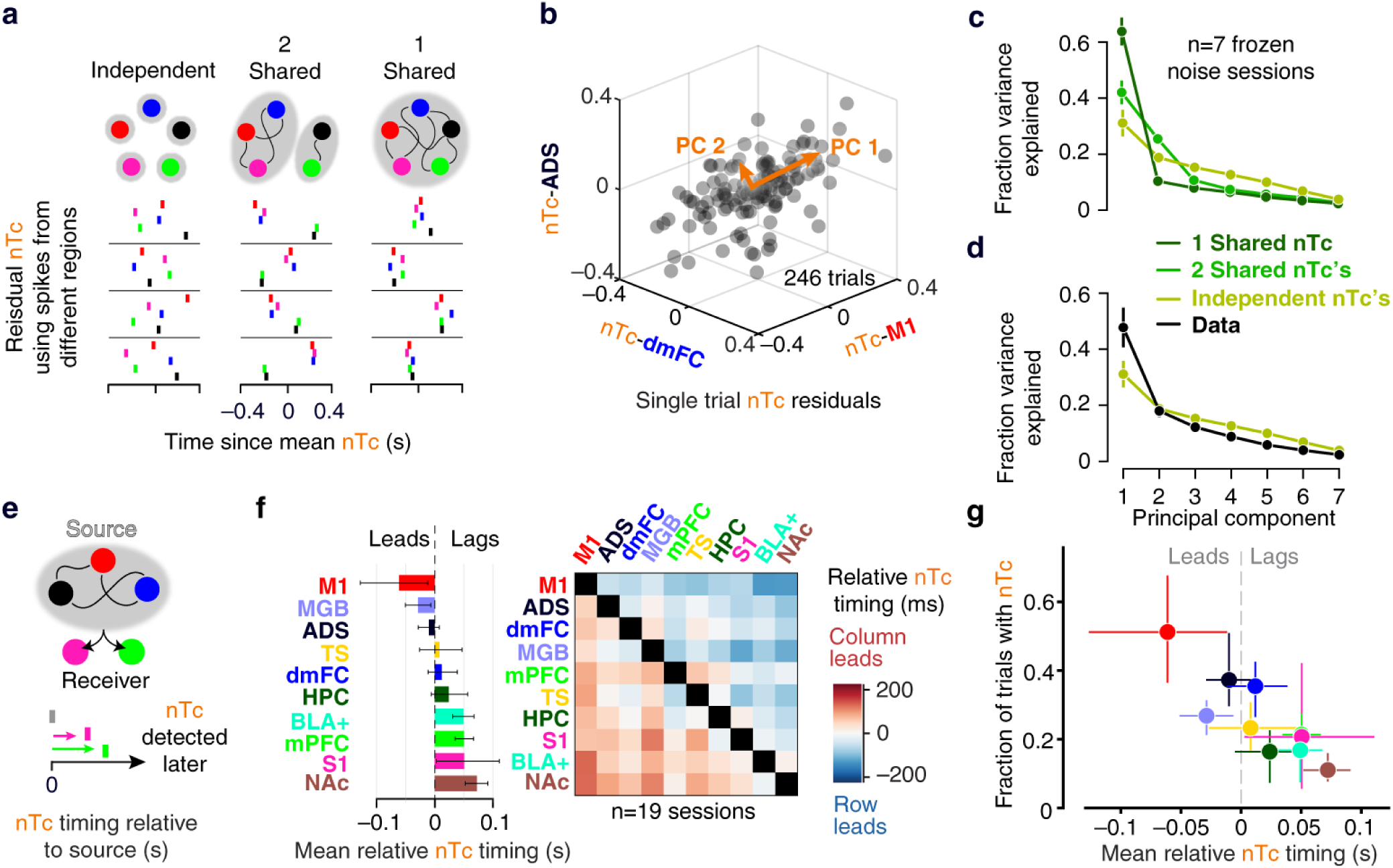
A single moment of covert decision commitment, led by M1, is shared across the brain. **a**, Schematic of competing hypotheses: the timing of decision commitment may be independent across brain regions or reflect one or more shared processes. Rows show four example trials with the same click train and behavioral choice. Tick marks indicate single-trial residual nTc for each region, relative to the average nTc for trials with that click train and choice. **b**, State-space representation of the coordinated variability in single-trial nTc residuals across 3 brain regions, shown for one session with 246 trials that had nTc estimates for all three regions. Orange arrows indicate directions explaining the maximum variance. **c**, Scree plot produced from applying probabilistic PCA to the set of across-region nTc residuals, generated from simulations implementing the competing hypotheses in (a) (see **Methods**). Error bars indicate 1 s.e. using a hierarchical bootstrap across sessions. **d**, Same as (c) but using the observed data. The prediction assuming independence across regions is reproduced for comparison, as it provides a measure of the noise floor. Error bars indicate 1 s.e. using a hierarchical bootstrap across sessions**. e**, Schematic of the approach for identifying where decision commitment signals first emerge. **f**, Left, mean relative nTc timing. Error bars are the bootstrapped 95% CIs over session-level trial-count-weighted means. Right, Matrix representing the trial-averaged difference in nTc timing between all pairs of brain regions. Choice AUC distributions were matched before calculating nTc (see **Methods**). Left, each region’s average difference relative to all others, corresponding to the mean of each row in the matrix on the right. Each pairwise comparison includes only trials with nTc detected in both regions. Negative values indicate that the row region’s nTc tends to lead the column region’s nTc. **g**, Scatter plot of the mean relative nTc timing from (f) the fraction of trials in which nTc could be detected from each region, computed on the choice AUC matched populations. **f-g**, Error bars are the hierarchically bootstrapped 95% CIs.

For the experimental data, because nTc was not detected on every trial from each region, we used probabilistic PCA (PPCA) to assess dimensionality in the presence of missing data (see **Methods**). We found that one dimension explained approximately 50% of the variance of the residual nTc’s, and the rest of the dimensions explained a fraction of variance that was below the noise floor (**Figure 6d**). This suggests a single, shared, nTc moment, with noise in its per-region estimate (and as examined immediately below, relative lags between regions). While this was most consistent with the MMDDM simulations that assumed a single, shared moment of decision commitment, we do note that the dominance of the first principal component here is not as strong as for the DV(t)s in **Figure 3i**. This may be due to the fact that nTc provides only one data point per trial, whereas the DV(t) residuals provide whole trajectories of many points per trial, implying less statistical power in our ability to distinguish a 1-dimensional nTc system from a 2-dimensional nTc system. We note that for trials with a given fixed stimulus, the only cross-trial difference in nTc estimates will be due to cross-trial differences in neural activity. In other words, the observation that nTc residuals are correlated is an observation about correlated firing patterns. In particular, the observation of a single dominant principal component suggests that changes in direction within each region’s neural activity space tend to happen at similar times across all regions.

We then measured lead-lag relationships between nTc’s from different regions (**Figure 6e**). To avoid confounds due to differences across regions in the strength of their choice encoding, for each pair of regions, we first subselected neurons from the each region’s recorded population, chosen until both regions had matched choice encoding strengths (**Methods**). nTc times were then estimated, for each region, from these matched data, and using those trials in which nTc was detected from both regions, we calculated the average difference in nTc timing between the two regions in the pair. These pairwise measurements were then summarized for each region by calculating the average nTc timing difference of that region with all others (**Figure 6f**). As with the finding that nTc could be detected most frequently from M1 (**Figure 5d**), this timing analysis revealed that decision commitment could be first detected from M1, with nTc-M1 leading nTc obtained from all other regions by an average of ∼50 ms (**Figure 6f**, left; per-rat results shown in **Figure S14**). Despite having first controlled for the strength of choice encoding, there was a strong correlation across regions (Pearson’s *r*=0.93) between nTc detection frequency and nTc timing (**Figure 6g**). In sum, analyses of nTc timing across regions reveal a low dimensional, highly coordinated representation consistent with a single, shared decision commitment process that can be detected earliest and most robustly in M1.

### Neural accumulation of evidence abruptly terminates at covert decision commitment

Here, simultaneous estimates of DV(t)s and nTc provided the opportunity to align each trial’s data on the putative time of covert commitment, and distinguish between the two classes of models, by obtaining a high resolution view of the timescale and the nature of changes in DV(t) dynamics around the covert commitment event. Since nTc and DV(t) are both estimated from noisy neural data, we avoided artefactual findings of a relationship between them by using disjoint sets of neurons to estimate each. For each region, we estimated DV(t) using neurons from that region, and estimated nTc using neurons from all other recorded regions (see **Figure S11d** for behavioral validation of leave-one-region-out nTc).

As in many previous studies^16,37,73^, DV(t)-derived choice prediction accuracy, averaged over trials and aligned either on stimulus onset (**Figure 3d**) or the subject’s initiation of the overt motor report (**Figure 7c**), ramped steadily upwards over time. Aligning trials on nTc instead revealed a dramatically different pattern: for almost all regions, choice prediction accuracy ramped upwards until nTc but then abruptly plateaued (**Figure 7d**; the two exceptions are MGB, where it falls instead of plateauing, and NAc, where it continues growing). This held even for trials in which nTc occurred many hundreds of milliseconds before movement onset, and also held when assessed through the unsigned DV(t) (**Methods**), which unlike choice prediction accuracy is not bounded at 1. We quantified the effect through bilinear regression on each of the unsigned DV(t) traces in **Figure 7e,f**. (For clarity, only the M1 bilinear fits are overlaid on the data here; see **Figure S15** for all such fits, including alignment to different timepoints and per rat). Aligned to movement onset, the slopes of both segments in the bilinear fits were positive, across all regions (**Figure 7g**). In contrast, when aligned to nTc, the slope of the first line segment was significantly positive for all regions (p<0.05 using hierarchical bootstrap test across sessions) while none were significantly positive for the second line segment (p>0.05 using hierarchical bootstrap test across sessions; **Figure 7h**).

**Figure 7.**
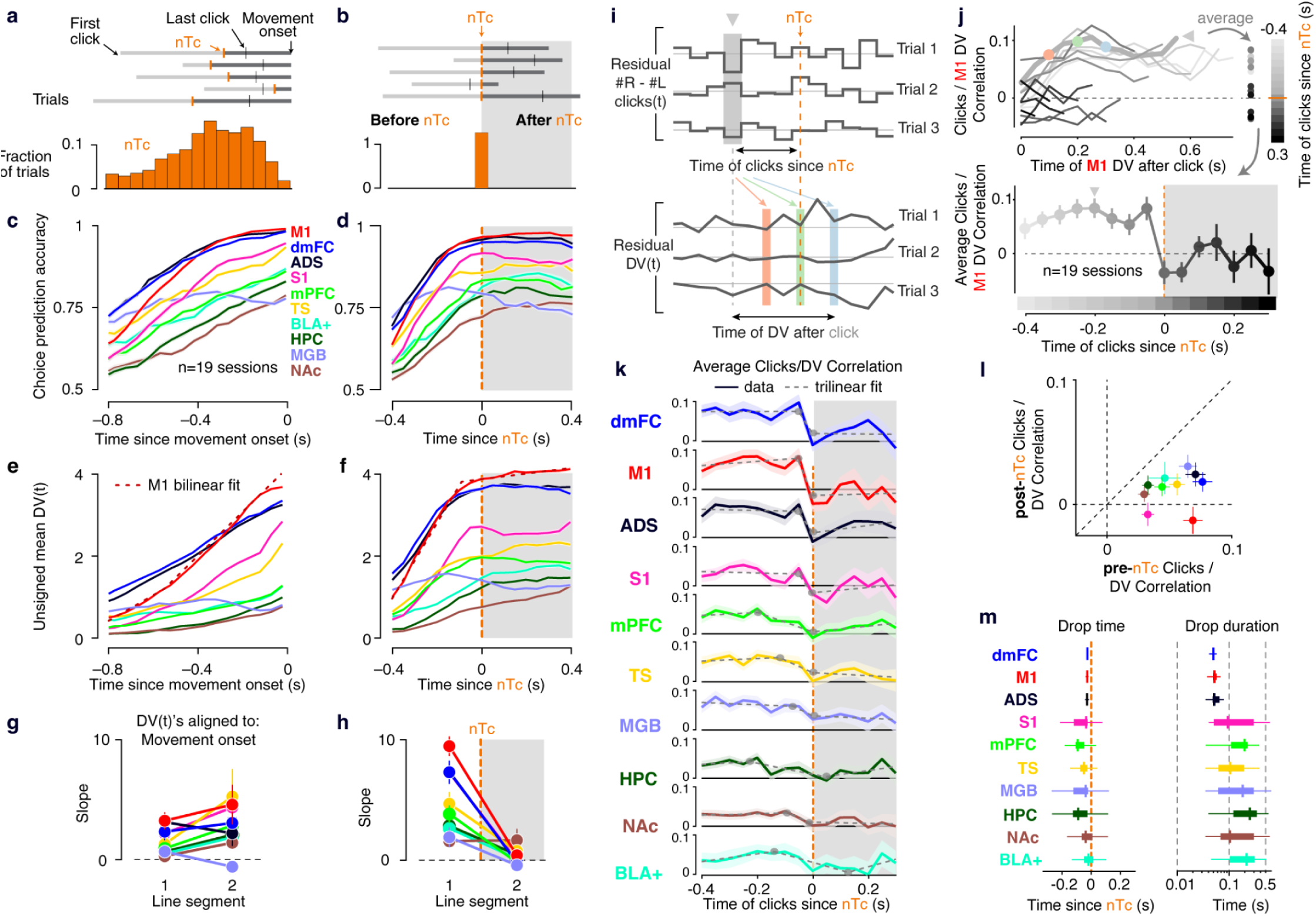
Neural accumulation of evidence abruptly terminates across the brain at covert decision commitment. **a**, Timing of nTc-M1 relative to movement onset. Top, sequence of the first click, nTc-M1, last click and movement onset for 5 example trials. Bottom, distribution of nTc-M1 across trials aligned to movement onset. **b**, Same as (a) with trials aligned to nTc-M1. **c**, Choice prediction accuracy using each brain region aligned to movement onset. Mean across 19 sessions shown. **d**, Same as (c), with trials aligned to nTc. Each region’s data was aligned to an estimate of nTc excluding spikes from that region, to avoid circularity. **e,f**, Trial-averaged DV(t) from each brain region, aligned to movement onset in (e) and each region’s respective “leave-one-region-out” nTc in (f). The sign of the DV(t)s on left choice trials was flipped before averaging. **g,h**, Slopes of the first and second line segments of bilinear fits of the mean DV(t) traces in (e) and (f), respectively. **i**. Correlation-based method to assess the influence of momentary sensory evidence on future evolution of DV(t)s. Briefly, trials are aligned to nTc and for each timebin, we calculate the across-trial correlation between the sensory evidence (#R-#L clicks) in that bin and a given region’s DV at future timepoints. The expected value of the sensory evidence and DV, given the generative click rate (ɣ, see **Methods**), is first subtracted to remove spurious temporal correlations. **j**. (top) Correlation between sensory evidence and future M1 DV(t) (result of applying method in i). Each curve reflects the influence of sensory evidence at a particular timepoint on future DVs. The example timepoint illustrated in (i) is highlighted as the bold gray curve. Dots in red, green and blue correspond to different future DV times indicated in (i). (bottom) Averaging the curves above gives a single time-varying correlation value capturing the sensitivity of DV(t) to clicks presented at different times. **k**. Same as bottom of, j but for all regions. In addition to the data (solid lines +/−1 bootstrap s.e.), a trilinear fit is also shown (dashed gray lines) with the two breakpoints indicated (gray circles). **l**. Average clicks-on-DV influence for clicks presented in the pre-nTc period versus in the post-nTc period. Error bars reflect +/−1 bootstrap s.e. **m**. Whisker plots showing drop times and drop durations across regions, obtained using the trilinear fits in (k). Drop time is defined as the midpoint of the second line segment and drop duration as the temporal duration of the second line segment. The distribution reflects fits to a trial-bootstrap distribution of the data.

This plateauing of trial-averaged choice prediction accuracy could be due to a loss in sensitivity of the DV(t)s to incoming sensory evidence at nTc. To directly probe this, we obtained a measure of incoming sensory evidence in 50ms bins relative to nTc on each trial and assessed how this influenced future values of DV(t). For each bin, we calculated the difference between the number of right and left clicks (#R-#L clicks), and subtracted the mean timecourse of #R-#L clicks across trials with the same generative left and right click rates (e.g., all trials with left clicks generated at 30 clicks/sec, right at 10 clicks/sec), yielding “residual #R-#L clicks” that were independent across time bins. Using the same groupings, we computed corresponding DV(t) residuals.

Then, for #R-#L residuals at each given time point relative to nTc, we measured the correlation coefficient between those #R-#L residuals and the DV(t) residuals at the same and future timepoints (**Figure 7i**). **Figure 7j** shows the results of this analysis for M1 DV(t)s. Sensory evidence presented before nTc (here using an nTc estimate that excluded M1 neurons) had an effect on future M1 DV(t) that took up to 0.2s to fully ramp up, and was then stable for hundreds of milliseconds, as expected from non-leaky accumulation with a sensory delay (lighter gray curves in **Figure 7j**). In contrast, as soon as the evidence was presented at nTc or later (darker gray curves in **Figure 7j**), the correlation between click residuals and M1 DV(t) residuals fell to zero: M1 DV(t) became insensitive to the clicks. We summarized each of the M1 curves in **Figure 7j** by its average, and in the red trace of **Figure 7k**, plotted those averages as a function of the time at which the evidence was presented. Other traces in **Figure 7k** are the corresponding analysis for the other recorded regions. For all regions, including those that were more weakly choice-selective, and even for sensory regions like MGB, the timecourse showed a decay aligned on nTc: clicks presented after nTc had a markedly reduced influence on DV(t)s compared to those presented before (**Figure 7l**, average reduction across regions of 75+/−8% s.e.). For the regions with the strongest choice-related activity (M1, ADS and dmFC), this reduction was remarkably abrupt and step-like at nTc. We quantified the reduction dynamics with a trilinear fit: the first segment captures the pre-nTc baseline, the second segment captures the drop, and the third segment captures the post-nTc steady state. The midpoint of the second segment provides a measure of the time of the drop time and its temporal extent quantifies the drop duration. For M1, ADS and dmFC, the drop time occurred less than one timebin before nTc and its duration was ≤50 ms, effectively instantaneous given the temporal resolution of the analysis (**Figure 7m**). For other regions with weaker choice-related activity (and therefore noisier DV(t)s), the trilinear fit exhibited greater uncertainty, making it difficult to make strong conclusions about the timing of the reduction. Nonetheless, and taken together with the widespread plateauing of DV(t)s at nTc, these results demonstrate an abrupt termination of evidence accumulation at covert commitment, strongly favoring models in which the transition from accumulation to commitment is a discrete, abrupt, and widely represented change in neural state – even in the absence of any overt action at that time.

## Discussion

Although it is well established that representations of accumulating evidence can be found in a widespread set of regions, the functional circuits underlying these widespread representations remain unresolved. And because covert decision commitment, by definition, occurs at a time that is hidden from view, its neurophysiological properties have been difficult to study. To address these issues, we applied and further developed two new tools. First, a recently-described method to estimate each trial’s timing of covert decision commitment^50^, and second, very large-scale multi-region simultaneous electrophysiological recordings, with which we recorded up to 4,032 units simultaneously across up to 20 regions covering a broad set of diverse regions of the brain.

### A single evidence accumulation process is shared across the brain

Models of evidence accumulation for decision-making have variedly proposed that accumulation occurs through a single accumulator, or through multiple distinct accumulators (for different evidence sources^5,29^, for different decision options^20–27^, or using distinct accumulation properties^28^). All of these models are consistent with evidence accumulation being represented broadly across the brain. To distinguish between these possibilities, we examined patterns of residual DV(t) co-fluctuations. While DV(t)s have been measured across multiple regions during short-term memory and for decisions involving simple binary stimuli^36,37,74^ this is the first study recording DV(t)s simultaneously from more than two regions during gradual evidence accumulation. By measuring DV(t)s across sensory, association, and motor regions, in both hemispheres, and in both cortical and subcortical regions, our simultaneous recordings provided a unique opportunity to examine the large-scale pattern of DV(t) residual co-fluctuations during the evidence accumulation period. Remarkably, despite the number and variety of recorded regions, co-fluctuations across all regions were dominated by a single principal component. In other words, a single scalar sufficed to describe the full set of residual DV(t) co-fluctuations, across both hemispheres (**Figure 3h,i**). This conclusion held regardless of whether regions were divided into left versus right hemispheres or left-choice- versus right-choice-preferring neurons (**Figure S9**). These observations constitute significant neurophysiological evidence in favor of single-accumulator models of decision-making. In addition, the cross-region correlations that we measured will provide strong constraints on multi-region models of decision-making.

### A frontal subcircuit is the source of diffusion noise in the accumulator

Many mathematical psychology models of decision-making account for variability in subjects’ responses by assuming that the subjects’ evidence accumulator also accumulates noise, leading to diffusion in the accumulator. Despite decades of research using these models, the anatomical source of the diffusion noise has remained unaddressed. Our findings that DV(t) residuals have a strong signature of a diffusion process, in that their variance grows linearly with time^39,68^ for up to 800 milliseconds into the stimulus presentation (**Figures 3e, S5**), and that frontal region fluctuations in DV(t) residuals temporally lead all other regions (**Figure 4**), now suggest frontal regions as the anatomical source of the accumulated noise (i.e., diffusion). This opens the door towards studying the neural mechanisms that underlie the diffusion process. If the diffusion noise and the accumulated evidence are jointly expressed in a single accumulator, as assumed in standard models, and as is consistent with the diffusion-like properties we found in DV(t)s (which are themselves defined in terms of the animal’s choices), then the findings also suggest that the accumulator’s evidence signal is computed in frontal regions, to be then broadcast to the rest of the brain.

### A single neural moment of covert decision commitment is shared across the brain and led by M1

Due to covert commitment’s hidden nature, the cellular resolution neural dynamics that underlie it have been understudied. Applying the nTc estimator of covert commitment timing^50^ to neural activity data from every region we recorded, we found that even while nTc was detected in a greater fraction of trials in frontal regions compared to posterior regions, nTc could be detected, and validated, remarkably broadly. nTc could be detected even in canonically sensory regions like the medial geniculate body. Thus covert commitment, like evidence accumulation, is represented very broadly in the brain. Analysis of patterns of co-fluctuations in nTc residuals suggested a single covert commitment process that is shared across the brain (**Figure 6**).

M1 stood out as the region from which nTc could be detected earliest, and on the greatest proportion of trials. This was the case even though we controlled for different strengths of choice signals in different regions. Therefore, at least among the regions recorded in this study, M1 is the best candidate for the source of the covert commitment event. The leading role of M1 in commitment contrasted with the decision formation phase, during which a larger set of regions were found to have roughly temporally coincident DV(t) fluctuations (**Figure 4c**), and suggests that temporal relationships across regions may change over different phases of decision-making. Previous results with overt (not covert) decision commitment have linked overt commitment to motor regions and the gating of action initiation^4,75–77^, a finding that seems natural given the immediate temporal proximity of overt commitment and a motor action. Here, using covert commitment instead, we show that the relationship between commitment and motor regions may be even more profound: when commitment and the overt response are temporally separated by hundreds of milliseconds, M1 strongly and rapidly encodes the cognitive event of committing to a choice.

### Neural accumulation of evidence abruptly terminates at the time of covert decision commitment

Decades-old yet still competing classes of models disagree about the nature of the transition from accumulation to covert commitment, proposing either an abrupt or a continuous transition (**Figure 1c**). The nTc estimate of each individual trial’s moment of covert commitment gave us the opportunity to directly compare DV(t) dynamics before versus after nTc. We found evidence for a remarkably abrupt (≤ 50 ms, which was our measurement resolution) change in sensitivity of DV(t)s to incoming evidence at the time of nTc. This was particularly clear in frontal regions (**Figure 7**). These data are striking new evidence that strongly favor models with abrupt, not continuous, transitions to covert commitment. The transition occurs at a moment that varies substantially over trials and that could only be detected using large-scale cellular-resolution recordings.

We conclude that the division of decision making into two discrete distinct states, initially a conceptual formalism in behavioral models, is also instantiated in the brain as a rapid (≤ 50 ms), purely internal state change that is evident globally and occurs at the time of covert decision commitment. Knowing the timing of this change on individual trials will be an important tool in separating fundamentally distinct stages of decision making.

## Overall conclusions

Ultimately, an adequate account of how a decision unfolds in the brain must go beyond statistical accounts of decision making behavior towards mechanistic models that describe how many regions across the brain interact. Our results provide important new constraints on such an understanding. We show that the widespread presence of decision-related activity across the brain, observed in past work, does not reflect the presence of parallel decision making modules, but rather a system that evolves coherently on single trials. Further, neither gradual evidence accumulation nor the abrupt moment of covert commitment emerge from distributed interactions across the entire brain, instead emerging in specific subcircuits which we delineate here, and then propagating broadly. Finally, our data highlight the importance of using large-scale cellular-resolution recordings to detect internal signals and internal events, such as covert commitment, that are not timelocked to external stimuli or overt motor acts but are nevertheless critical features that fundamentally shape cognition’s neural dynamics and their relation to behavior.

### Limitations of the study

In this study we performed unusually widespread simultaneous cellular-resolution recordings across the rat forebrain. However, many regions were not recorded in this study, notably including the entirety of the midbrain, hindbrain and cerebellum, all of which have been linked to decision making. We cannot rule out the possibility that recordings from these regions would alter some of the conclusions made based on the present data. Notably, the superior colliculus (SC), which was not recorded here, is the region most closely linked to overt decision commitment in non-human primates^76,77^. Relating SC’s overt commitment signatures with the results reported here is an important topic for future study.

Our results do not reveal what triggers the abrupt and brain-wide neural change at the time of covert decision commitment. In many decision making models, the level of evidence reaching a predetermined bound is the trigger, but the results here provide no direct evidence in favor or against this. At least in the context of covert commitment, it is plausible that other neural processes, unrelated to the level of accumulated evidence, could be responsible for triggering this change.

## Acknowledgments

We thank Zoe Boundy-Singer, Bing Brunton, Mark Goldman, Ben Lankow, Maxwell Melin, Hendrikje Nienborg, Bence Olveczky, Maxwell Shinn, and Corey Ziemba for their helpful suggestions and comments. We also thank Jessica Morrison, Klaus Osorio, Jovanna Teran, Andres Bustos and Emily Valance for technical assistance. We thank Grace Barnett and Jamus MacGuire for veterinary advice. We are grateful to all members of the Brody lab for their support, collegiality and feedback.

This manuscript was supported in part by the National Institutes of Health (NIH) under grants R01MH108358, R01MH138935, and 5U19NS132720. It is subject to the NIH Public Access Policy. Through acceptance of this federal funding, NIH has been given a right to make this manuscript publicly available in PubMed Central upon the Official Date of Publication, as defined by NIH. Additional support was provided by the Howard Hughes Medical Institute Investigator Program.

## Author Contributions

**Conceptualization**, A.G.B., J.A.C., T.Z.L., T.H., and C.D.B.

**Methodology**. A.G.B., J.A.C., and T.Z.L.

**Formal analysis**. A.G.B. led the majority of data analysis, with contributions from T.Z.L., C.D.B, J.A.C, K.S., and E.Y.X.

**Investigation**. A.G.B., J.A.C., T.Z.L., C.D.K., W.S., S.J.V., L.L., and S.O.

**Data Curation**. A.G.B., J.A.C., S.J., and T.Z.L.

**Visualization**. A.G.B., J.A.C., T.Z.L., W.S., and M.B.

**Writing - Original Draft**. A.G.B., J.A.C., and T.Z.L.

**Writing - Reviewing & Editing**. A.G.B., J.A.C.,T.Z.L., C.D.K., W.S., S.J.V., and C.D.B.

**Supervision**. C.D.B.

**Project administration**. A.G.B., J.A.C.,T.Z.L., and C.D.B.

**Funding acquisition**. T.H. and C.B.

## Data Availability

The datasets generated during the current study are available from the corresponding authors upon reasonable request and will be deposited in a public repository prior to publication.

## Code Availability

All code used in the analyses will be made freely available upon publication.

## Methods

### Subjects

Three adult male Long-Evans rats (Hilltop) were used for the experiments presented in this study. All procedures were approved by the Princeton University Institutional Animal Care and Use Committee and were carried out in accordance with National Institutes of Health standards. Rats were pair-housed in Technoplast cages until their implantation surgery and kept in a 12 hour reversed light-dark cycle. All training and testing procedures were performed during the rat’s dark cycle. Rats had restricted access to water such that the water consumed daily was at least 3% of their body mass.

### Behavioral task

Rats performed the behavioral task in custom-made training enclosures (Island Motion, NY) within sound- and light-attenuated chambers (IAC Acoustics, Naperville, IL). Each enclosure consisted of three straight walls and one curved wall in which three nose ports were embedded (one in the center and one on each side). Each nose port contained one light-emitting diode (LED) as well as an infrared (IR) beam to detect the entrance of the rat’s nose into the port. A loudspeaker was mounted above each of the side ports and was used to present auditory stimuli. Each of the side ports also contained a small metal tube that delivered water reward, with the amount of water controlled by valve opening time.

Rats performed an auditory discrimination task in which optimal performance required the gradual accumulation of auditory clicks^41^. At the start of each trial, rats inserted their nose in the central port and maintained this placement for 1.5 s (“fixation period”). After a variable delay of 0.5-1.3 s, two trains of randomly timed auditory clicks were presented simultaneously, one from the left and one from the right speaker. Regardless of onset time, the click trains terminated at the end of the fixation period, resulting in stimuli whose duration varied from 0.2-1 s. The train of clicks from each speaker was generated pseudo-randomly by an underlying Poisson process, with different mean rates for each side. The combined mean click rate was fixed at 40 Hz, and trial difficulty was manipulated by varying the ratio of the generative click rate between the two sides. The generative click rate ratio (*γ*) varied from 39:1 clicks/s (easiest) to 20:20 (most difficult). At the end of the fixation period, rats could orient towards the nose port on the side where more clicks were played and obtain a water reward.

Psychometric functions were computed by dividing trials into eight similarly sized groups according to the total difference in the right and left clicks, and for each group, computing the fraction of trials ending in a right choice. The confidence interval of the fraction of right response was computed using the Clopper-Pearson method. Recording sessions were excluded if the rat completed <300 trials or showed a lapse rate (λ) > 8%, estimated using the following psychometric model:

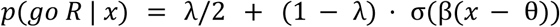

where *x* is the total difference in the number of right and left clicks, θ is the threshold (i.e., point of subjective equality), β is the sensitivity (i.e., steepness of the curve), and σ(·) is the logistic (sigmoid) function.

Two rats consistently maintained low lapse rates (i.e., the error rate on the easiest trials), each with a median lapse rate of 2% and maximum of 5% in each session, and completed a large number of trials (median 623 per session; range: 370-806). The third rat exhibited greater variability across sessions; we excluded from analysis 4 of 14 sessions with lapse >8% and 2 sessions with fewer than 300 trials (see **Figure S3**).

### Frozen noise

In the Poisson Clicks task, the click trains for each trial depends on that trial’s generative click rate ratio (*γ*) and a seed that determines the sample of two pseudorandom variables: the stimulus duration and the Poisson processes determining left and right click times. For most of the 19 sessions analyzed, stimulus seeds were unique for each trial. For a subset of seven sessions across the three animals (“frozen noise” sessions) a small number of unique stimulus seeds (9 for each of the 6 values of *γ* used) were interleaved throughout the session. Only these sessions contributed to the analyses involving DV(t) and nTc residuals. In the main text, this corresponds to Figs. 3f-j, Figure 4, and Figure 6b-d.

### Implant

Previous methods^78–81^ have focused on reusability, surrounding each probe with a bulky enclosure from which it could be removed at the end of the experiment. Here, we sought instead to maximize probe density and flexibility in target selection, forgoing reusability by directly cementing each probe to the skull. Only a small probe holder was used, to provide an attachment point for a micromanipulator used to lower the probes into the brain. This probe holder had two components: one that is glued permanently to a probe, and another that is removed after the probe is anchored to the skull, to reduce the volume and weight of the implant. The arrangement of the probes on the skulls of the rats was performed using CAD software, as was the design of the probe holder and a “chassis” that surrounded the probes and held the headstages (**Figure S3a-e**). The probe holders were 3D printed in-house using the Form 3 SLA printer (Formlabs) in Black V4 resin (Formlabs; RS-F2-GPBK-04) and the headstage holder was printed in Tough 1500 (Formlabs; RS-F2-TO15-01). CAD files for these components can be found at https://github.com/Brody-Lab/uberphys_paper/tree/main/CAD_files.

After printing the parts, they were visually inspected and sanded to ensure proper mating. The probe holder parts were secured using two 4 mm M1.2 screws (McMaster; 96817A746). The headstage holder was assembled using 3mm M1 screws (McMaster; 96817A704) and headstages were secured to the headstage holder using 4 mm M1.2 screws (McMaster; 96817A746). To secure the probes in the probe holders, each probe holder was placed in a stereotaxic cannula holder (Kopf, Tujunga, CA, USA; Model 1766-AP Cannula Holder) which was held in place by a vise. The probe was then placed on the probe holder and aligned to the axis of the cannula holder. A small amount of thick-viscosity cyanoacrylate glue (Mercury Adhesives) was applied to the edges of the probe holder using a small wooden dowel.

### Surgery

Surgery was performed using similar techniques to those reported previously^79^, but with several important innovations to support the lengthy and invasive nature of the 8-probe implantation. All surgical procedures were performed under isoflurane anesthesia (1.5-2%) using standard stereotaxic technique. Rats were given an intraperitoneal (IP) injection of ketamine (60 mg/kg), ketofen (5 mg/kg) and Ethiqa XR (0.65 mg/kg) to assist induction and provide analgesia. To ensure proper hydration throughout the surgery, rats were given 3 mL saline subcutaneously after induction and every 3 hours afterward.

The dorsal skull was exposed by making an incision along the rostral/caudal orientation along the top of the head. The skull surface between the lambdoid sutures and 20 mm anterior of the frontonasal suture was cleaned and scrubbed. The temporalis muscle was detached from the lateral ridge and retracted to gain access to the tail of the striatum. The sites of nine craniotomies, one for the ground cannula (Protech International, 22G/5mm), and eight for the Neuropixels 1.0 probes were marked with a sterile pen. The craniotomy for the ground had a diameter of approximately two millimeters, and each craniotomy for a Neuropixels 1.0 probe had a diameter of 1 mm. The 3D profile of each craniotomy intended for a Neuropixels 1.0 probes had a conical shape to minimize the amount of dura exposed (to maximize the stability of the chronic recording) while maximizing the range of angles through which the dura can be accessed, thereby facilitating the subsequent durotomy. After completing the nine craniotomies, they were covered with Gelfoam (Pfizer), and then dental cement (C&B Metabond Quick Adhesive Cement System) was applied to the skull surface. Durotomies were made using a 27G needle and fine forceps. After the ground cannula was lowered, the craniotomy was sealed with a silicone adhesive (KWIK-SIL, World Precision Instrument). The cannula was adhered to the skull through a dental composite (Absolute Dentin, Parkell).

Neuropixels probes were stereotaxically inserted into the brain using a motorized micromanipulator (Narshige, MDS-1) at a speed of ∼5 µm/s. Each craniotomy in which a probe was inserted was sealed using a silicone gel (Dowsil 3-4680), applied using a micropipette. The Neuropixels probes are bonded to the skull and existing fixtures using dental composite.

After all eight probes had been inserted, the silver wire shorting the ground and reference pad of each probe were twisted together and soldered onto the ground cannula. To reinforce the attachment between the probes and the skull, liquid dental acrylic was applied to the skull surface. To shield the probes and to mount the headstages, a chassis (**Figure S3**) was attached to the fixtures using dental composite.

The surgeries were considerably longer than typical rat intracranial implant surgeries, lasting >12 hours. A team of five surgeons contributed in rotating shifts. We also provided the rats extended analgesia for two days post-op and ad libitum water access for at least one week after surgery.

### Electrophysiological recording

Neural activity was recorded using chronically implanted Neuropixels 1.0 probes that were permanently affixed to the skull using custom-designed 3D-printed probe holders described above. We used acquisition hardware from NI (a PXIe-1071 chassis) in conjunction with SpikeGLX software (https://github.com/billkarsh/SpikeGLX) to acquire the data. The reference selected for each probe was a silver wire shorted to the ground wire and penetrating the olfactory bulb. The amplifier gain used during recording was 500. Spikes were sorted offline using Kilosort2^82^, using default parameters and without manual curation. In each of three animals, probes bilaterally targeted one of four locations described in detail in Table 1. We excluded 3 sessions from the rat A327 due to an error in specifying the electrodes to be recorded.

**Table 1.** Recording targets. Four insertion targets were used, bilaterally in each subject. Probes were sometimes angled in either or both the sagittal and coronal plane, both to accommodate multiple probes on the subject’s head and to target specific combinations of brain regions. A positive angle in the sagittal plane indicates that the probe tip was more anterior than the probe base. A positive angle in the coronal plane indicates the probe tip was more lateral than the probe base. To avoid blood vessels and collisions between probes, some variability in coordinates across subjects was required. In these cases, the range of coordinates is indicated.

| Site # | 1 | 2 | 3 | 4 |
| --- | --- | --- | --- | --- |
| Craniotomy coordinates, mm relative to Bregma (AP, ML) | +4.2, 1.0 | +1.9, 3.0 | [-2.0 : -2.1], [5.0 : 5.2] | [-5.7 : -6.0], 3.7 |
| Insertion angle in sagittal plane (deg) | 0 | 0 | [0 : 5] | [0 : 5] |
| Insertion angle in coronal plane (deg) | -10 | -10 | 5 | 0 |
| Insertion Depth (mm) | [3.9 : 4.9] | [7.4 : 7.9] | [6.8 : 7.6] | [7.4 : 7.9] |
| Regions targeted | dmFC, mPFC | M1, ADS, NAc | S1, TS, GP, BLA+, Pir | V1, HPC, DS, MGB, SBN, SN |

### Histology

Rats were transcardially perfused with 10% formalin under anesthesia with 0.4 mL ketamine (100 mg/ml) and 0.2 mL xylazine (100 mg/ml) IP. Brains were cleared using modified uDisco, volumetrically imaged using lightsheet microscopy, and aligned to the Princeton RAtlas^51^. These methods are described in detail elsewhere^51^.

While the Princeton RAtlas provides a useful tool for visualizing brains in a common coordinate space, adding well validated region annotations to the RAtlas remains a work in progress. Therefore, to assign recorded units to brain regions, we used the following procedure. Using the BigDataViewer^83^ Plugin for Fiji^84^, we dynamically resliced the lightsheet volumes to obtain virtual slices that best visualized each individual probe track. These virtual slices were then segmented into brain regions by visual comparison to the Paxinos and Watson rat atlas. The recording sites on each probe were then assigned to a position within the virtual slice, by converting from image pixels to physical distance given the insertion depth of each probe. We found we could more accurately estimate the insertion depth of each probe from electrophysiology rather than by using the nominal insertion depth recorded during surgery. The electrophysiological estimate was determined by the most superficial channel on each probe at which multi-unit activity could no longer be clearly observed.

### Neuronal selection

Units were only included for analysis if they exceeded predefined thresholds for a number of quality metrics based on waveform shape. These thresholds are defined in Table 2 below, and were designed to exclude units that were a) not of biological origin, i.e. noise artifacts; and 2) not of somatic origin, since axonal spikes could be generated by fibers of passage. These criteria are highly similar to those recently proposed by another group to exclude artifacts and non-somatic units from silicon probe recordings^85^. In addition, for decoding analyses and estimation of decision variables, units were only included if they fired at least 1 spike on at least half of trials (i.e. whose “presence ratio” exceeded 0.5). Approximately 65% of units found by Kilosort2 were included given these criteria. Note that no criteria were applied to exclude multi-units. Additionally, because multiple sessions were recorded from the same animals, the same neurons might have been sampled across days.

**Table 2.** Waveform-shape-based unit inclusion criteria. Where applicable, metrics are defined for the average waveform on the main channel (i.e. the channel for which the unit had the largest peak-to-peak voltage).

| *Quality metric* | *Description* | *Allowed values* |
| --- | --- | --- |
| Spatial spread | Spatial decay constant of an exponential fit to the waveform energy as a function of distance to the peak site | <150 μm |
| Peak width | Width of main deflection at half height | <1 ms |
| Peak-trough width | Time from trough to peak | <1 ms |
| Upward-going spike | Has an upward-going peak deflection | FALSE |
| uVpp | Peak-to-peak voltage | >50 μV |

### Neural decoding of choice and calculation of “decision variables”

We used logistic regression to decode choice from population firing rates. Neuronal firing rates were obtained by convolving spike times with a 50 ms s.d. symmetric Gaussian smoothing filter and sampled at 50 ms intervals. A separate logistic regression model was fit to each 50 ms sample. The model probability of a rightward choice at a given time point *t* on trial *i* was *p_t_*_,*i*_ (*R*) = *f*(*X_t_*_,*i*_ β*_t_* + α*_t_*), where *f* is the sigmoidal logistic function, *X_t_*_,*i*_ is the vector of neuronal firing rates for time point *t* on trial *i*, β*_t_* is the vector of weights for time point *t*, and α*_t_* is a model constant for time point *t*. The subject’s choice on trial *i* is modeled as *C_i_* ∼ *Bernoulli*(*p_t_*_,*i*_ (*R*)), where *C* = 1 corresponds to a rightward choice. β*_t_* and α*_t_* are the model parameters, shared across trials and chosen to maximize the log-likelihood of the subject’s choices: i.e. *argmax* Σ*_i_ log p*(*C_i_* |*p_t_*_,*i*_ (*R*)). We used 10-fold stratified cross-validation within sessions to assess model performance as well as to identify the optimal L1 regularization hyperparameter for β*_t_*. The “decision variable” for a given time point and trial *DV_t_*_,*i*_ was defined as the linear predictor *X_t_*_,*i*_ β*_t_* + α*_t_*, which is equivalent to the log-odds of *p_t_*_,*i*_ (*R*), i.e. *log* (*p_t_*_,*i*_ (*R*)/*p_t_*_,*i*_ (*L*)):

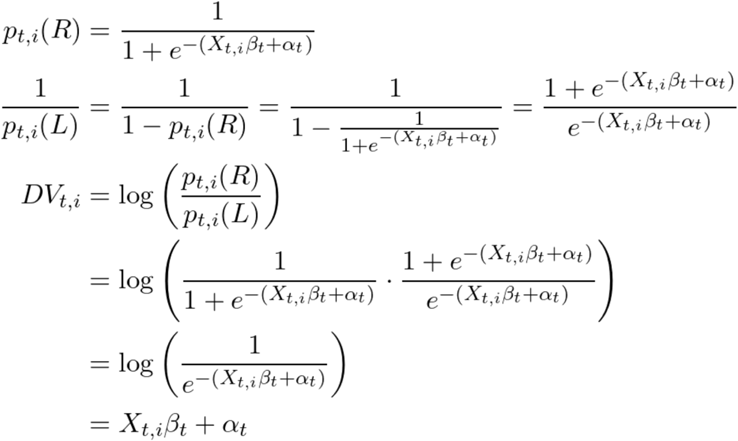

To ease comparisons of DV(t)s across timepoints within a trial, we used a constant value of the L1 regularization hyperparameter across timepoints, obtained as the geometric mean of the hyperparameters identified through cross-validation at each timepoint. Choice prediction accuracy was calculated for each session using the “balanced accuracy”. The standard or *unbalanced* choice prediction accuracy (*CPA*) across a set of *n* trials for time point *t* can be given as:

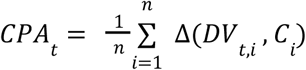

where Δ is an indicator function that equals 1 if *sign*(*DV_t_*_,*i*_) = *C_i_* and 0 otherwise. The balanced accuracy is the average of the unbalanced accuracy computed separately for left and right choice trials. This removes upward bias associated with an uneven distribution of choices. To fit the decoding models, we relied on the Glmnet^86^ package in MATLAB R2024a (Mathworks, MA, USA). A separate decoding model (and associated choice prediction accuracy curve and set of DV(t)s) was performed for each region. When DV(t)s and choice prediction accuracy curves are shown without separating by hemisphere, a decoding model that included neurons in both hemispheres was used (as opposed to using per-hemisphere models and averaging the results). Only timepoints between the first click on each trial and the moment the animal began its choice reporting movement (the moment it broke fixation and pulled out of the center port) were used for choice decoding.

### Neural decoding of momentary evidence and calculation of “momentary evidence variables”

We used linear regression to decode momentary evidence from population firing rates (**Figure S8**), using a very similar procedure to decoding of choice. Only the differences are stated here. Neuronal firing rates were obtained by convolving spike times with a 50 ms s.d. symmetric Gaussian smoothing filter and sampled at 100 ms intervals. Momentary evidence was defined as the difference in right versus left clicks within a 100 ms window before the time of the firing rate was sampled. A separate regression model was fit to each 100 ms sample. The model prediction of momentary evidence at a given time point *t* on trial *i* was *e_t_*_,*i*_ (*R*) = *X_t_*_,*i*_ β*_t_* + α*_t_*, and we refer to this as the “momentary evidence variable” (MEV), and is analogous to the DV in the choice decoding model. It is a 1D summary of the neural population’s estimate of the momentary stimulus evidence. The model parameters were chosen to maximize the squared error of the momentary evidence estimates. As with choice decoding, 10-fold stratified cross-validation within sessions was used to assess model performance as well as to identify the optimal L1 regularization hyperparameter for β. To fit the decoding models, we again relied on the Glmnet^86^ package in MATLAB R2024a (Mathworks, MA, USA).

### Calculating DV(t) Residual (Cross)-Correlations

To estimate the component of the DV(t)s that reflected noise in the evidence accumulation process itself, as opposed to signals related to movement preparation or stimulus encoding, we subtracted from a trial’s DV(t) trace the mean across trials with the same frozen noise seed and choice.

DV(t) residual correlations were calculated as the Pearson correlation between the DV(t) residuals estimated from different brain regions, concatenating across trials and time points. A single correlation value was calculated for each pair of regions on each of the 7 frozen noise sessions and a single grand mean was obtained for each pair of regions, averaging across sessions. To focus on the time period during which the subject was actively accumulating evidence, only timepoints up to each trial’s neurally estimated time of commitment (nTc-all; using all recorded neurons) were used, and trials with no detected nTc-all were excluded. The same procedure was also used to calculate correlations between other projections of neural activity, including “momentary evidence variables”, as in Figure S8.

In Figure 4, DV(t) residual correlations were also calculated as a function of different lags between pairs of regions (i.e. DV(t) residual cross-correlograms). Again a separate cross-correlogram was calculated for each pair and session, and then the mean across sessions was obtained. The peak lag in this grand-mean cross-correlogram was calculated as the lag that produced the maximum value within the range [-0.3,0.3]s. To calculate the mean peak lag of a given region relative to all others, we averaged the peak lags of that region and all others.

To assess uncertainty of the mean peak lag, we repeated the entire procedure described in the previous paragraph 500 times, each time resampling trials with replacement from the set of 7 frozen noise sessions, to create a bootstrap distribution of mean peak lags for each region.

### Assessing Dimensionality of DV(t) Residuals

We used principal components analysis (PCA) to assess the dimensionality of the DV(t) residuals across regions. To do this, we define a matrix *X* where each column corresponds to the DV(t) residual for a single region in each hemisphere (i.e., a feature) and each row corresponds to a single time point in a trial (i.e., a sample). We concatenated time points across trials within a session, and centered the data. PCA transforms the coordinates of *X* such that the axes are ordered by maximum variance explained. The principal component scores (i.e. the coordinates of the original data in the transformed space) are given by *T* = *XW*. The columns of *W* are equivalent to the eigenvectors of *X^T^X*. The variance of *X* explained by the *K^t^*^ℎ^ principal component is equal to λ*_k_*/ Σ*_k_* λ, where λ*_k_* is the *k^t^*^ℎ^ eigenvalue of *X^T^X*. The loadings *L* are given by the columns of *W* scaled by the square root of the corresponding eigenvalues (i.e. *L* = *W*Λ, where Λ is the diagonal matrix of eigenvalues). Elements of *L* give the cross-covariance of the activity of each region with its projection onto each PC, providing a measure of pairwise alignment between the regions and the PCs. To focus on the time period during which the subject was actively accumulating evidence, only timepoints up to each trial’s neurally estimated time of commitment (nTc-all; using all recorded neurons) were used, and trials with no detected nTc-all were excluded.

To estimate uncertainty around the results of PCA, we performed a hierarchical bootstrap across sessions as follows. We performed the above procedure for PCA for each session 1000 times, each time resampling trials within a session with replacement. For each resample, a statistic for a given PC (e.g. loading for a given region, fraction variance explained) is obtained for each session and then averaged across sessions. The hierarchical nature of the bootstrap is achieved by, instead of averaging across all sessions, resampling with replacement from the set of 7 frozen noise sessions and then averaging the statistic across those resampled sessions.

To generate predictions for the PCA dimensionality analysis under assumptions of different numbers of latent decision processes, we used a set of shuffling procedures. To generate predictions for a single latent decision process, we shuffled the anatomical identity of each cell. This creates “pseudo-regions” of the same size as the original populations recorded in each region, but containing cells randomly sampled from all recorded regions. We then obtained DV(t) residuals from these pseudo-regions, and performed PCA as described above. This shuffling procedure tests the idea that the decision-related activity in each region are i.i.d samples from the same underlying distribution.

To generate predictions for the PCA results assuming complete independence across regions, we randomly shuffled time points of *X* separately for each region (columns of *X*). To generate predictions assuming two independent decision processes, we first randomly divided regions (columns of *X*) into two groups and randomly shuffled time points of each group. In this way, temporal alignment was abolished between groups but preserved within groups. These shuffling procedures were applied many times to estimate shuffle-based distributions which were then further sampled as part of the hierarchical bootstrap across sessions used to generate confidence intervals in the plots shown in Figure 3.

### Relating DV(t) Residuals to Spontaneous Movements

We considered the possibility that DV(t) residuals, and correlations between these measures across regions, reflects shared encoding of spontaneous movements. To assess these, we assessed how well the DV(t) residuals could be predicted using overhead behavioral video recorded during each recording session. Video was recorded at 25Hz. We nonlinearly compressed each video into a small set of latent variables using a convolutional autoencoder, following the encoder/decoder architecture adapted from BehaveNet^87,88^. Each frame was converted to grayscale, resized to 128 × 128 pixels, and scaled to the range [0, 1]. The encoder passed each frame through four convolutional layers (32, 64, 256, and 512 channels; 5 × 5 kernels; stride 2, so the image was halved at each layer; leaky-ReLU nonlinearities), followed by a fully connected projection to a 16-dimensional bottleneck; a mirror-image decoder of transposed-convolutional layers reconstructed the frame, ending in a sigmoid nonlinearity. We trained one autoencoder per session to minimize the mean-squared reconstruction error between each input frame *x* and its reconstruction *x*^,

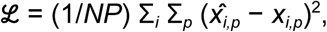

summed over the *P* pixels of each of the *N* frames in the batch, using the Adam optimizer (learning rate 10^−4^) for 200,000 weight-update iterations, each on a random minibatch of 900 frames. After training, the encoder mapped every frame to its 16-dimensional latent vector. We rotated the latents onto their principal components and whitened them, so that the resulting video latents were mutually uncorrelated and of unit variance.

We then assessed how well the 16 video latents for each session could predict a given region’s DV(t) and DV(t) residual. Because DV(t)s were sampled at 50ms intervals relative to the first click, while video was recorded at 25Hz using a different clock, we first interpolated the set of video latents using shapewise-preserving cubic interpolation at the times of the DV(t) samples. Then, a separate linear regression model was fit to each 50 ms timestep relative to the first click. The optimal L1 penalty was obtained separately for each timestep under 10-fold cross-validation. We did this both for the raw DV(t)s as well as the DV(t) residuals, computed after subtracting the mean within groups of trials with the same stimulus seed and choice. The variance explained was calculated separately for each timestep as well as jointly across all timesteps. To estimate the uncertainty of variance explained, we used a hierarchical bootstrap across sessions and trials. To do this, for each of many resamples, we first randomly sampled (with replacement) from the set of 6 frozen noise sessions with video. Then, for the sampled set of sessions, we sampled (again with replacement) from the pooled set of trials across those sessions. Then variance explained for that bootstrap sample was calculated across that set of trials.

### Linear network models

We modeled each brain region as a population of *n* = 100 units and described the activity of the full system, comprising *R* = 18 regions, by a state vector **x**(*t*) ∈ ℝ*^nR^*. Activity evolved under linear, normal (orthogonally diagonalizable) dynamics,

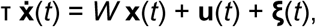

where *W* is the connectivity matrix, τ = 100 ms is the dynamical time constant, **u**(*t*) is the stimulus drive (which were unit positive impulses for right clicks, unit negative impulses for left clicks, slightly smoothed in time by a gamma function with 25 ms time to peak), and **ξ**(*t*) is noise. Equation 1 was integrated with the Euler–Maruyama method (10 ms steps) from the onset of the click stream to the end of the trial. Inputs were the same for all line attractors in the model, and were always parallel to the line attractors.

The connectivity matrix was constructed so that each functionally independent sub-network embedded a single line attractor. For a sub-network spanning a set of ρ regions, we began in a ρ-dimensional space with a diagonal dynamics matrix whose eigenvalues were 0 along one integration direction (the line attractor) and −1 along the remaining, stable directions. We then rotated this space so that the line attractor pointed along a direction with unequal weights across the dimensions ρ, each of which represented one region; the evidence input therefore moved different regions by different amounts, endowing each region with its own decision-variable (DV) gain. This rotation did not change the nature of the dynamics, but did change the connectivity matrix for the ρ regions so it now had non-zeros on the off-diagonals. Each region was then expanded from one to *n* dimensions by appending *n* − 1 additional stable directions (eigenvalue = −1), with no coupling among these extra *n*−1 units within a region or across regions. At this point the weight matrix for the sub-network was now *n*ρ-dimensional, with only ρ(ρ−1) non-zero off-diagonals (due to the line attractor, defined across regions), and (*n*^2^−1)ρ^2^ − (*n*−1)ρ off-diagonals that were zero (all the units that are uncoupled to each other). However, we then applied an independent random orthogonal rotation within each region’s *n*-dimensional subspace. These within-region rotations leave the eigenvalues, and hence the dynamics, entirely unchanged. But they render the weight matrix for the subnetwork dense and all-to-all even though communication between regions in the subnetwork remained rank one. Different subnetworks were all independently built in this manner.

To couple different subnetworks to each other (since otherwise they were entirely uncoupled), we defined an additional mode before the region-wise rotation. Defined jointly across the second unit in every region of all sub-networks in the full model, this mode, which had dimensionality *R* and whose connectivity matrix was initially diagonal, had eigenvalues drawn uniformly from [−5, −0.5]. Since they are all negative, any noise in this mode decays away. We then randomly rotated the mode’s *R*-dimensional space. This created an additional *R*^2^−*R* non-zero off-diagonals in the full connectivity matrix (which is much larger, with *n*^2^*R*^2^ elements). The non-zero off-diagonals introduced by this mode included non-zero off-diagonals across different sub-networks. Since the line attractors were defined in the first unit of each region, and this mode was defined in the second unit, the two remained entirely separate and did not affect each other. When each region was randomly rotated within its own *n*-dimensional space, this now rendered the full connectivity matrix *W* dense with all-to-all connectivity, even though the underlying dynamics were simple normal dynamics with entirely independent line attractors, defined across separate dimensions.

The stimulus drive was derived from the click trains presented to the animals. We convolved each right-and left-click time with a causal, unit-area gamma kernel (shape parameter 2; time to peak, 25 ms) to obtain a smooth momentary drift rate, and drove the line attractor of each sub-network with the difference between the right and left convolved trains. Because the kernel integrates to unity, each click contributed a fixed increment to the long-time DV(t). At every integration step we added independent Gaussian noise of standard deviation σ√(*dt*) to all dimensions (σ = 4); this diffusion was the only source of trial-to-trial variability in the DV(t)s.

We read out the DV(t) of each region as the projection of its population activity onto the region’s line-attractor direction, estimated from a random subset of *m* = 30 of its *n* = 100 units. Reading out fewer units than the network contains introduces a readout noise that lowers the correlations among regional DV(t) residuals while leaving the predicted choice — the sign of the DV, which is dominated by the full-dimensional signal — essentially unaffected. To match the processing done with the experimental data, regional DV(t)s were smoothed in time with a Gaussian kernel (standard deviation, 50 ms) before further analysis.

We simulated three architectures with identical anatomy (18 regions of 100 units) that differed only in how the regions were partitioned into functionally independent sub-networks: a single sub-network spanning all 18 regions, two sub-networks spanning 9 regions each, and 18 sub-networks each confined to a single region. We then analyzed DV(*t*) residuals from the model in the same way as done with DV(*t*) residuals in the experimental data.

### Variance of DV(t) residuals versus time

For each brain region we used frozen noise sessions, and took the held-out (cross-validated) population decision variable DV(*t*) residuals around each random seed’s mean trajectory, as a function of time *t* relative to the first click. We restricted the analysis to the pre-commitment epoch: on trials with a detected time of neural commitment (nTc) we discarded all time bins later than *t*_nTc_ − 0.2 s, while trials with no detected commitment contributed their entire trajectory. Pooling these residuals across seeds, we computed the across-trial variance at each time bin, Var(*t*), from the first click onward (*t* ≥ 0).

We fit the per-bin variance with the linear model

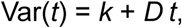

by weighted least squares, weighting each bin by the number of contributing residuals minus one. Here *k* is the time-zero (readout) variance and *D* the diffusion rate of the underlying accumulation. Fits were computed separately for every (session, region) pair; for the across-region summary, the per-bin variance traces were averaged over the seven frozen-noise sessions before fitting. To compare the form of the relation across regions on a common scale, each trace was rescaled by its own fitted parameters, Var(*t*) ↦ (Var(*t*) − *k*)/*D*, so that an exactly linear trace falls on the line *y* = *t*.

We quantified the quality of the linear fit by the weighted coefficient of determination, *R*^2^, computed for each (session, region) pair. Per-bin variances and their uncertainties were estimated treating the frozen-noise seed as the independent unit — the across-seed mean of the within-seed variance at each bin, with its standard error from the seed-to-seed scatter — and we confirmed these uncertainties with a seed-level bootstrap (200 resamples). We report *R*^2^ for each region in every session, with a bin required to draw on at least three seeds, each contributing at least six trials, to enter the fit.

### Multi-mode drift-diffusion model (MMDDM)

A detailed description of the multi-mode drift-diffusion model (MMDDM) can be found in Ref.^50^. Briefly, MMDDM consists of a dynamic model governing the time evolution of a 1-dimensional latent variable and measurement models specifying the conditional distributions of the observations (spike counts and behavioral choice) given the value of the latent variable.

In the dynamic model, when the value of the latent variable (*z*) is not at either absorbing bound *-B* or *B*, its value at each time step t depends on the momentary input (*u*), which is corrupted by multiplicative noise of variance *σ_s_*^2^, and additive noise ε:

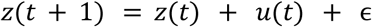

When *z* reaches either bound, it remains at the bound. The dynamic model has three free parameters, the bound height *B=(10, 20)*, variance of the multiplicative noise *σ_s_*^2^*=(0.1, 20)* and the mean of the initial state of the latent variable *μ_0_=(−5, 5)*. We chose to fit the input-related noise (rather than other sources of noise) because previous work suggests it to be the dominant source of noise in our task^41^. The additive noise on each time step is an i.i.d Gaussian with variance *Δt*, which is the time step *Δt=0.01* s.

The measurement model of the behavioral choice depends on only the sign of the latent variable *z* on the last time step of each trial (positive indicating rightward). There is no free parameter in the measurement model of the choice.

The measurement model of the spike count of neuron *n* at time step *t* is given by

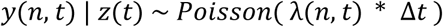

The firing rate λ, which has the unit of spikes/s, is

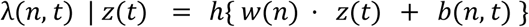

where *h* is the softplus rectifying nonlinearity, *w* the neuron’s encoding weight of *z*, and *b* a time-varying baseline input that is independent of *z*, the left or right clicks, or the animal choice. The baseline *b* accounts for time-varying influences of neural activity aligned to two events in the trial, stimulus onset and movement onset and slow drifts over minutes across a session. The measurement model of each neuron has 19 free parameters.

The parameters of the dynamic model and the measurement models are learned simultaneously. The gradient of the log-likelihood of the model has a closed-form expression and is used to optimize the parameters using the L-BFGS algorithm. Only responsive (> 2 hz) and choice-selective neurons (among the subset selected under the criteria described in “Neuronal selection”) were included in the fitting of MMDDM. Choice selectivity was computed using an ideal observer analysis, the receiver operating characteristic (ROC), categorizing between left and right choices using the spike counts during the first 0.5 s from the stimulus onset on each trial (excluding trials ending before 0.5 after stimulus onset). Choice selectivity is defined as |area under the ROC - 0.5| > 2^−5^ (median choice selectivity among responsive neurons=0.0315). For each session, a single model was fit to all neurons combined across brain regions, rather than fitting separate models to individual regions. Using the model parameters (θ) optimized from this joint fit, we then computed separate posterior probability distributions based on spikes from individual regions:

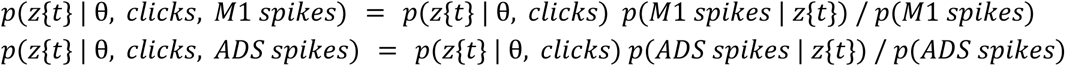

Each region-specific posterior distribution depends on the evidence provided by the spikes from that region alone. The posterior was computed using spikes both before and after that time step (i.e., it is the smoothed, not filtered, posterior).

After fitting MMDDM, we simulated spike trains by drawing samples from the latent process on the same set of trials used to fit the model. Each simulated trial used the original click train of the corresponding real trial. For each trial, we began by computing the prior distribution over the latent state at the first time step, *p*(*z*{1}), and drew a sample 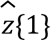 from this distribution. We then computed the conditional prior for the next time step, 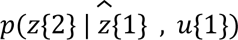), where *u*{1} is the click input at time 1, and drew a sample 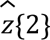. This process was repeated sequentially to generate a full sample path 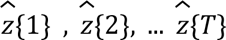 from the latent process for the *T* time steps in the trial. Given this latent trajectory, we computed the firing rate of neuron *n* on time *t* is computed as 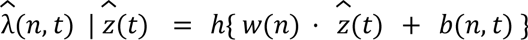 and drew a spike count from a Poisson distribution with rate 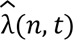.

### Neurally inferred time of commitment (nTc)

The time of decision commitment is defined as the first moment when the posterior probability of the latent variable in MMDDM being either bound exceeds 0.95 and remains above 0.95 for the remainder of the trial.

### Psychophysical kernel aligned to nTc

The psychophysical kernel quantifies the influence of momentary evidence on the animal’s upcoming choice, as a function of time relative to nTc. The kernel was estimated using a reverse correlation approach. To calculate the momentary evidence contained in the auditory clicks stimulus, we took the difference between the number of right and left clicks in each trial *i* for each 50-ms bin *t* (Δ*clicks*(*i*, *t*)) relative to that trial’s nTc. Because the generative click rate ratio (*γ*) varied across trials, this introduces temporal correlations in Δ*clicks* which reduces the estimation efficiency of reverse correlation. To ameliorate this, we subtracted the expected value from each trial, and assessed how the residual fluctuations in Δ*clicks* correlated with choice. We coded the animal’s choices either 1 (go-Right) or 0 (go-Left) and similarly demeaned them by subtracting the empirical average value of the choice across groups of trials with the same value of *γ*. The value of the psychophysical kernel at each time point *t* was then calculated as the across-trial Pearson correlation between the demeaned Δ*clicks*(*t*) and the animal’s demeaned choices. Only timepoints on each trial that occurred during the stimulus presentation were included in the analysis

### Assessing Dimensionality of nTc residuals

To calculate nTc residuals, for each region, we separated trials into groups with identical click trains (frozen noise seed) and behavioral choices, computed the mean nTc within each group, and subtracted it from the corresponding trials. We define a matrix *X* where each column corresponds to the nTc residuals for a single region in each hemisphere (i.e., a feature) and each row corresponds to a single trial (i.e., a sample). Because a value for nTc was not detected from every region on every trial, *X* has many missing elements. We therefore used probabilistic principal components analysis^89^ (PPCA) instead of PCA, for its ability to gracefully handle missing data. PPCA models the data as *X^T^* = *WT^T^* + ε, where *W* is a matrix of orthonormal coefficients (as in PCA), *T* are the latent factors (analogous to the scores in PCA), and *ε=v*∗*I*(*k*) is an isotropic error term that accounts for unexplained variability in the data. For *ε* to be non-zero, *T* must have rank *k* that is lower than the original data. Otherwise, PPCA simply reduces to regular PCA. Under the model specified by PPCA, *X* ∼ *N*(0, *W* * *W* + *v*∗*I*(*k*))). The parameters are estimated using EM. We used the PPCA function in MATLAB R2024a (Mathworks, MA, USA) to fit the model to the data. Principal component variances and loadings are computed in the same way as described above for PCA applied to the DV(t) residuals, except in this case we use the eigenvalues of the fitted model estimate of *X*.

To generate predictions for the dimensionality of the brain-wide nTc residuals under assumptions of different numbers of latent decision processes, we used simulations from MMDDM (see “Multi-mode drift-diffusion model (MMDDM)” in Methods) and applied the same procedure for computing nTc from the simulated spike trains as for the real data. Simulations were run on the same set of trials used to fit the model, preserving both the number of trials and the click train stimuli.

To model a single latent process, we simulated spike trains across all regions using a single run of the MMDDM. To model two latent processes, we simulated two independent runs for each trial and randomly assigned each brain region to use either the first or the second run consistently across all trials (i.e., a region assigned to the first run uses only latent trajectories from that run on every trial). To model fully independent latent processes for each region, MMDDM simulations were computationally impractical, since this would require simulating as many independent runs as brain regions. Instead, we used the same shuffling approach used for the DV(t) residuals, randomly shuffling time points of *X* separately for each region (columns of *X*).

### Quantification of plateau in decision-related activity at decision commitment

To quantify the rise and then abrupt plateau of the trial-average DV(t)s aligned to nTc-M1 (Figure 7C), we fit the trial-average DV(t)s for each region on each session using piecewise linear regression using two segments (i.e. a bilinear model). Specifically, a value of the mean DV(t) at time *t* was modeled as:

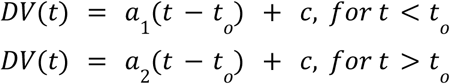

Where *_o_t* is the time of the breakpoint between the two line segments, *a*_1_ and *a*_2_ are the slopes of the two line segments and *c* is a model constant, in this formulation parameterized as the predicted value of *DV*(*t_o_*) (the mean DV(t) at the breakpoint). The four model parameters (*t_o_*, *a*_1_, *a*_2_, and *c*) were found that minimized the mean squared error, using the *lsqcurvefit* function in MATLAB R2024a (Mathworks, MA, USA).

### Calculating the Time-Varying Influence of Sensory Evidence on DV(t)s

To assess whether and how nTc affects the representation of accumulated evidence, we used a statistical approach similar to the method described above to calculate psychophysical kernels relative to nTc. Whereas the psychophysical kernel captures the influence of sensory evidence on choice as a function of time, this analysis seeks to capture the influence of sensory evidence on the DV(t)s. We calculated each trial’s Δ*clicks*(*t*) trace as described above, subtracting the expected value given the generative click rate ratio. For a given region, we demeaned each trial’s *DV*(*t*) trace in the same way. We only considered trials with detectable nTc and realigned both Δ*clicks*(*t*) and *DV*(*t*) traces to this time on individual trials, so that *t*=0 corresponds to nTc. To avoid potential circularity caused by a region contributing to the estimation of DV(t) and nTc, for each region we used a leave-one-region-out nTc in which that region’s spikes were not used in the estimation of nTc.

After realigning to nTc, We retained Δ*clicks*(*t*) over [-0.4,0.3]s and *DV*(*t*) over [-0.4,0.6]s. Trials were then pooled across all 19 sessions, and timepoints not overlapping with the stimulus period were excluded. We then calculated the full cross-correlation matrix of the demeaned Δ*clicks*(*t*) and *DV*(*t*) (i.e. the Pearson correlation between Δ*clicks*(*t*) and *DV*(*t* + τ) for all possible values of *t* and τ). Because a click can only influence the DV(t) at a later time, the informative entries lie below the τ=0 diagonal, and each diagonal corresponds to a fixed value of τ, i.e. time lag. A column of the lower-diagonal portion of the matrix captures the influence of Δ*clicks*(*t*) at a particular timepoint on future DV(t)s at all values of τ, and are what is shown in Figure 7j, top. Only correlation values to which at least 1200 trials contributed were included.

We then obtained a single correlation value at each timepoint relative to nTc (i.e. a single function of *t*) that captures the influence of sensory evidence on all future DV(t)s, regardless of τ, and is what is shown in Figure 7k for each region. This is equivalent to taking the mean of the columns in the lower-diagonal portion of the cross-correlation matrix. We did not simply average across all possible values of τ, because not every lag carries a meaningful click-on-DV signal due to sensory delays, the unique integration timescale of different regions, and limits in signal-to-noise at longer lags in which there are fewer trials with valid timepoints. To decide which lags to keep, the correlation is averaged along each diagonal (lag) using *only click times before nTc* — the epoch in which clicks are known to drive the DV. A lag is retained if its average pre-nTc click-on-DV correlation is significantly above chance, defined using the distribution of acausal correlation values (i.e. values in the cross-correlation matrix above the diagonal). This yields a region-specific set of significant lags.

As described in the main text, the function that captures the influence of sensory evidence at different times relative to nTc on future DV(t)s (i.e. the column means of the cross-correlation matrix described above) showed a step-like drop in clicks-on-DV influence at nTc for many regions. To quantify this step-like drop, we performed a trilinear fit, independently for each region and each of 300 resamples of the data, so that the fitted parameters and fitted curves inherit a bootstrap distribution. We used a hierarchical bootstrap across sessions and trials (cite). The trilinear model has six parameters and is the continuous piecewise-linear function:

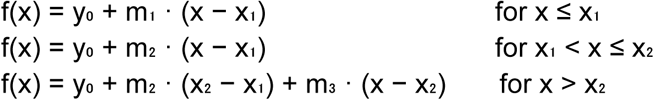

where m₁, m₂, m₃ are the slopes of the three line segments, x₁ < x₂ are the two breakpoints, and y₀ = f(x₁) is a vertical offset. Continuity across breakpoints is guaranteed by construction. We constrain x₁ to lie in the first half of the data range. All six parameters are fit simultaneously by nonlinear least squares (using “lsqcurvefit” in MATLAB). To guard against local minima, three initial guesses are tried — a thirds split of the time range, a narrow middle segment centred near t = 0, and a “flat-ramp-flat” configuration in which only the middle slope is non-zero — and the lowest-residual fit is retained. The final fit parameters were used as a warm-start initial guess (in addition to the automatic starts described above) when fitting each individual bootstrap resample.

### Choice AUC distribution matching

To ensure comparisons of nTc timing were not confounded by differences in overall choice selectivity, we computed for each pair of brain regions a set of units from each region whose choice-selectivity distributions are statistically matched. Choice selectivity was calculated for each unit as Receiver Operator Characteristic-Area Under the Curve (ROC-AUC) for leftward vs. rightward choices. First, ROC-AUC was computed on spike counts in a 500 ms window from stimulus onset. Trials with stimulus duration < 500 ms were excluded from this analysis. Then, units with |AUC - 0.5| ≤ 0.03125 and units with firing rates less than 2 spk/s were excluded from further analysis. For each of the brain region pairs, |AUC-0.5| distributions were matched independently within each recording session using the following algorithm:

1. Sort each region’s units by |AUC – 0.5| in ascending order.
2. Compute all combinations of possible trims of units from the high and low end of the above list for the reference region. Each trim direction (high and low) is capped at 50% of the region’s population size. Each of the regions takes a turn as the reference region.
3. For each possible trim, compute the mean and max values of |AUC – 0.5|.
4. For each trim of the reference region, search for the best trim of the non-reference region such that the following criteria are met:

a. The |AUC – 0.5| means are within 0.005
b. The |AUC – 0.5| maxes are within 0.005
c. The trims are within 10 units in population size
5. Select the trim pair that retains the most units among the valid trim pairs identified in step 4.

To compute pairwise lead-lag estimates of nTc timing, the resulting AUC-matched populations were fit with MMDDM and used to calculate the nTc of each region.

### CNN video classification and nTc decoding analysis

We used a SqueezeNet convolutional neural network (CNN) model^90^ to perform binary classification on extracted video frames and assess whether the model could distinguish between pre-nTc and post-nTc behavioral states. Video frames were extracted at specific temporal intervals (50, 100, 200, and 400 ms) relative to three target events: nTc, overt movement onset, and a temporally shuffled nTc control.

The rationale for temporal shuffle control is that video frames from later portions of a trial may be distinguishable from frames from earlier portions, even when the frames are not specifically aligned to nTc. The temporal shuffling process disrupts the trial-by-trial alignment between the actual nTc time and the extracted video frames. For each trial, a first-click-to-nTc latency is borrowed from another randomly selected trial and applied to the current trial’s first-click time. This generates a shuffled nTc time that preserves the overall distribution of first-click-to-nTc latencies while disrupting the timelocking to the true nTc event for that trial.

To prepare the data for CNN classification, we constructed composite three-channel images for each pre-event and post-event time point. Each composite image included the video frame closest to the target time, along with the immediately preceding and subsequent frames.

### Statistical tests

Unless otherwise stated, error bars and p-values are calculated non-parametrically, using bootstrap resampling either across trials or hierarchically across sessions. All p-values were computed using two-sided tests and not adjusted for multiple comparisons, unless otherwise specifically stated.

To test whether nTc-M1 was more likely or occurred earlier on trials with stronger evidence ([#R-#L]/[#R+#L]), we performed separate regression analyses restricted to the tertile of trials with longest stimulus durations. Logistic regression was used to model the binary outcome of whether nTc-M1 occurred on a given trial as a function of evidence strength, and for trials where it occurred, linear regression was used to model the latency of nTc-M1 as a function of evidence strength. Each model included only evidence strength and a constant term and no additional covariates.

