## Supplemental Information for "Multi-region coordination of evidence accumulation and decision commitment in the brain"

### The PDF file includes:

Supplementary Figures 1-15

Supplementary Table 1

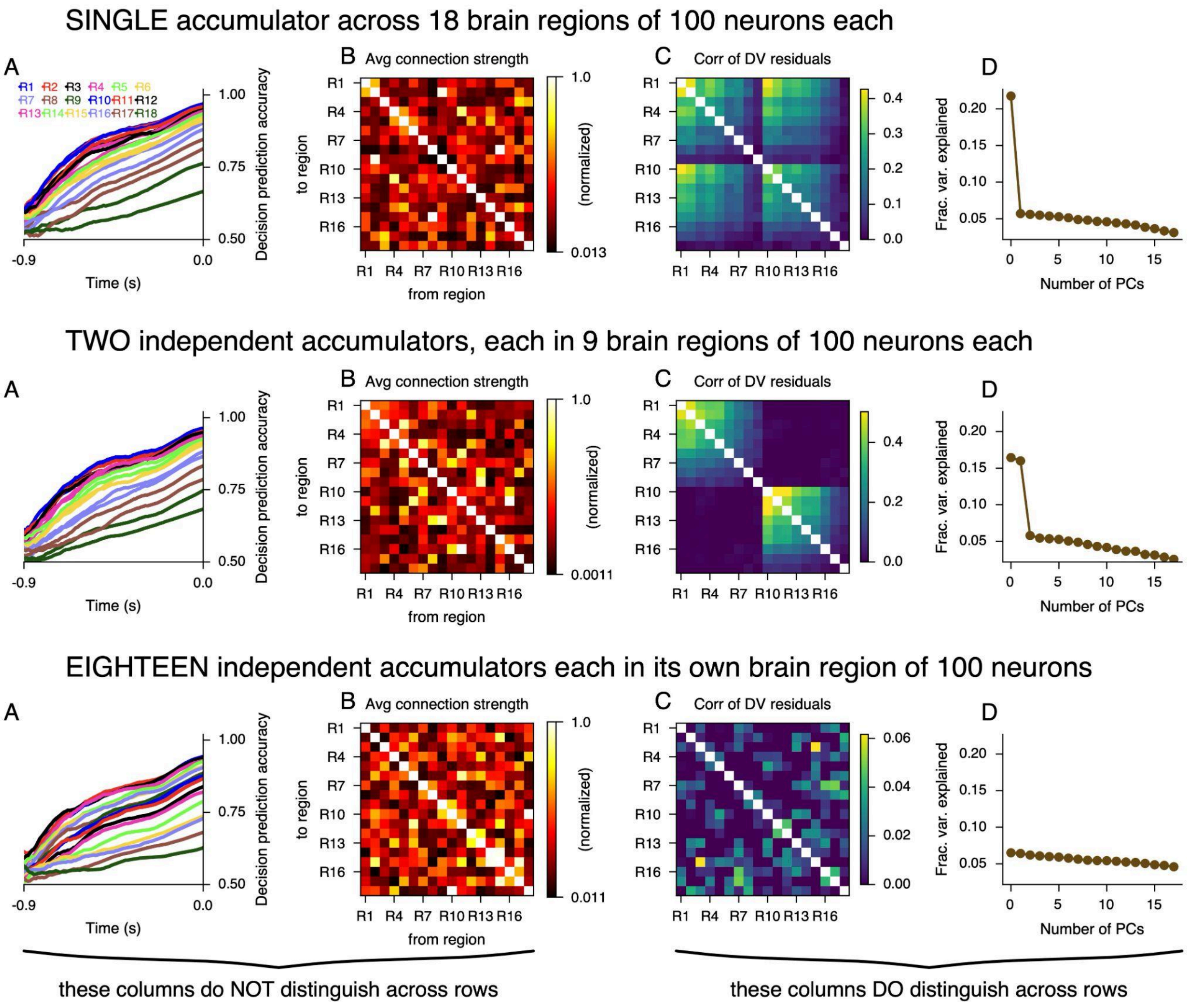


**Figure S1. Three synthetic models, all with similar representations across brain regions, but very different functional architectures, are distinguished by the pattern of co-fluctuations in their DV(t)s, related to Figures 1 and 3.** Each model had 18 brain regions with 100 neurons each, and had line attractor linear dynamics constructed to allow the network to accumulate sensory evidence. **A**) We computed DV(t) for each brain region, in response to the same sensory stimuli and trials as used in our experiments, and used each region's DV(t) to predict the probability of a correct decision choice. RX stands for "Region X". Across all models and brain regions, prediction accuracy rises over time as the models accumulate evidence. Compare to main text **Figure 3d**. **B**) In all three models, all regions are connected to all other regions. Random components in the model construction lead to different details in mesoscale connectivity, but all instantiations have non-zero connectivity between every pair of regions. **C**) Patterns of DV(t) co-fluctuations distinguish the three models from each other. Shown are correlation coefficients in DV(t) residuals around the trial means with a given stimulus, as in main text **Figure 3e**. **D**) Principal component spectra of the correlation matrices shown in (**C**) distinguish the three models, and indicate the number of separate functional circuits (here, number of line attractors) underlying the neural activity in each model. Compare to main text **Figure 3h.** Top row: a single evidence accumulator, shared across all regions of the model "brain". Second row: two independent accumulators, each housed in a distinct set of regions (in a real brain, these could correspond to left vs right hemispheres). Notice that mesoscale connectivity does not reveal the disjoint set of regions that correspond to each line attractor. Third row: one independent accumulator per brain region. DV(t) co-fluctuations are essential for distinguishing the models' different functional circuit architectures.

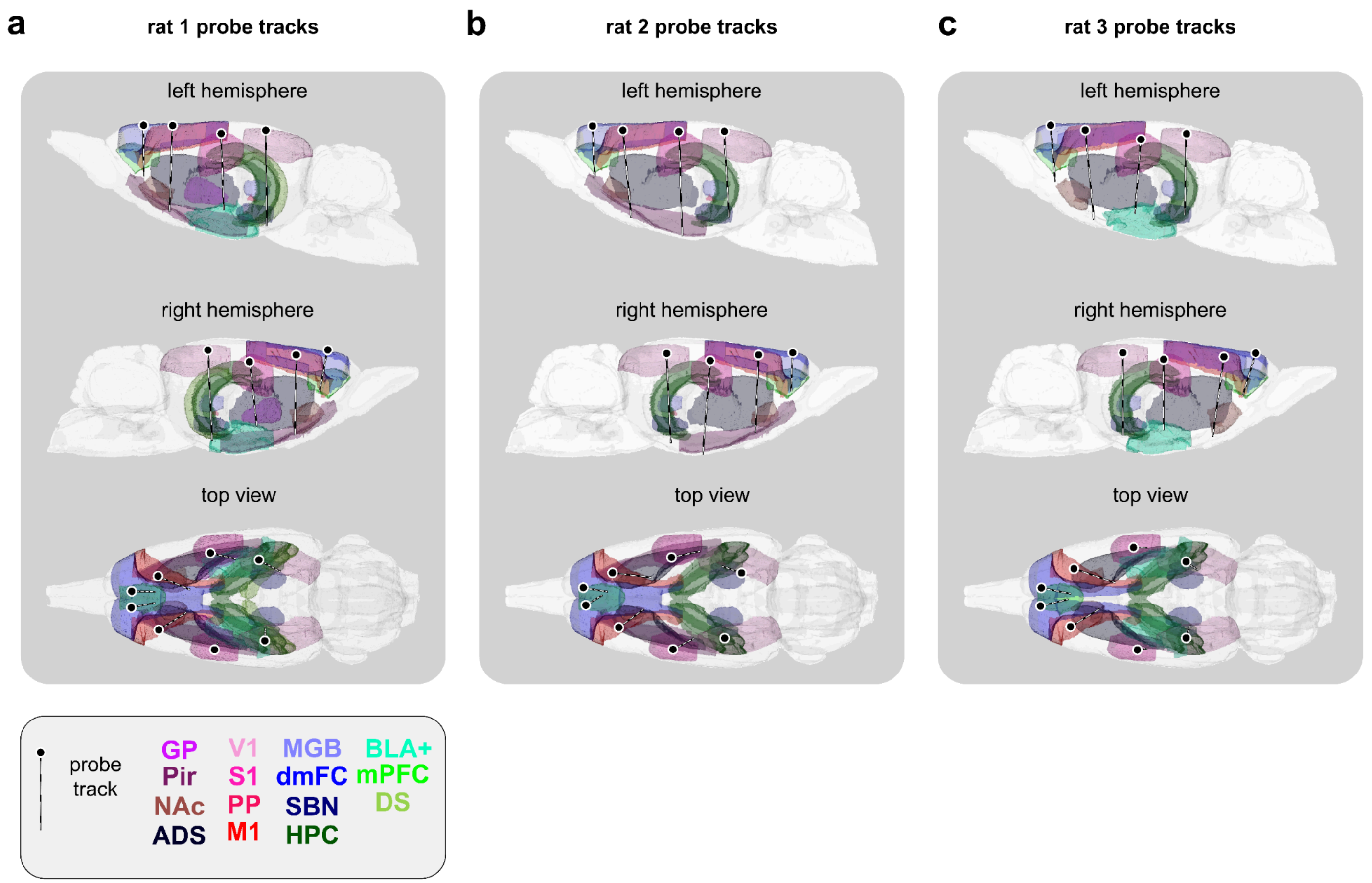


**Figure S2. Atlas registered brains for each individual rat, related to Figure 2. a**, The 8 implanted probe tracts of rat 1 reconstructed in the space of a common atlas volume. All regions we recorded units in rat 1 are outlined by annotations from the Waxholm Space rat atlas [15]. See Table S1 for full region names and lists for each animal. **b-c**, Same as **a** for rat 2 and 3, respectively.

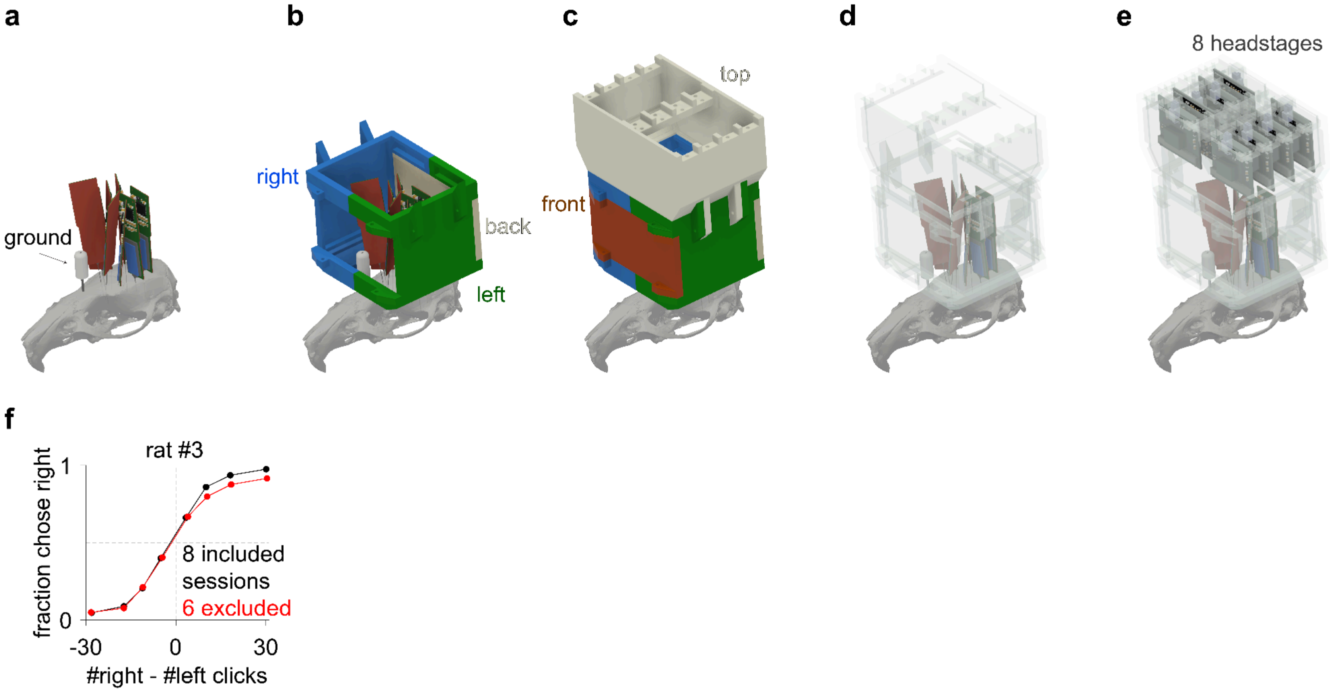


**Figure S3. Chassis for protecting the probes and mounting the headstages and behavioral performance of rat, related to Figure 2. a**, Eight Neuropixels 1.0 probes and a ground cannula. **b**, Three separate 3D-printed components that mate together to form the right, left, and back of the chassis. **c**, The front and top pieces. **d**, View of the probes and ground cannula within the chassis. **e**, The eight headstages are mounted on the top piece. **f**, For rat #3, four recording sessions were excluded on account of a visible lapse, and two excluded due to the number of trials completed being less than 300.

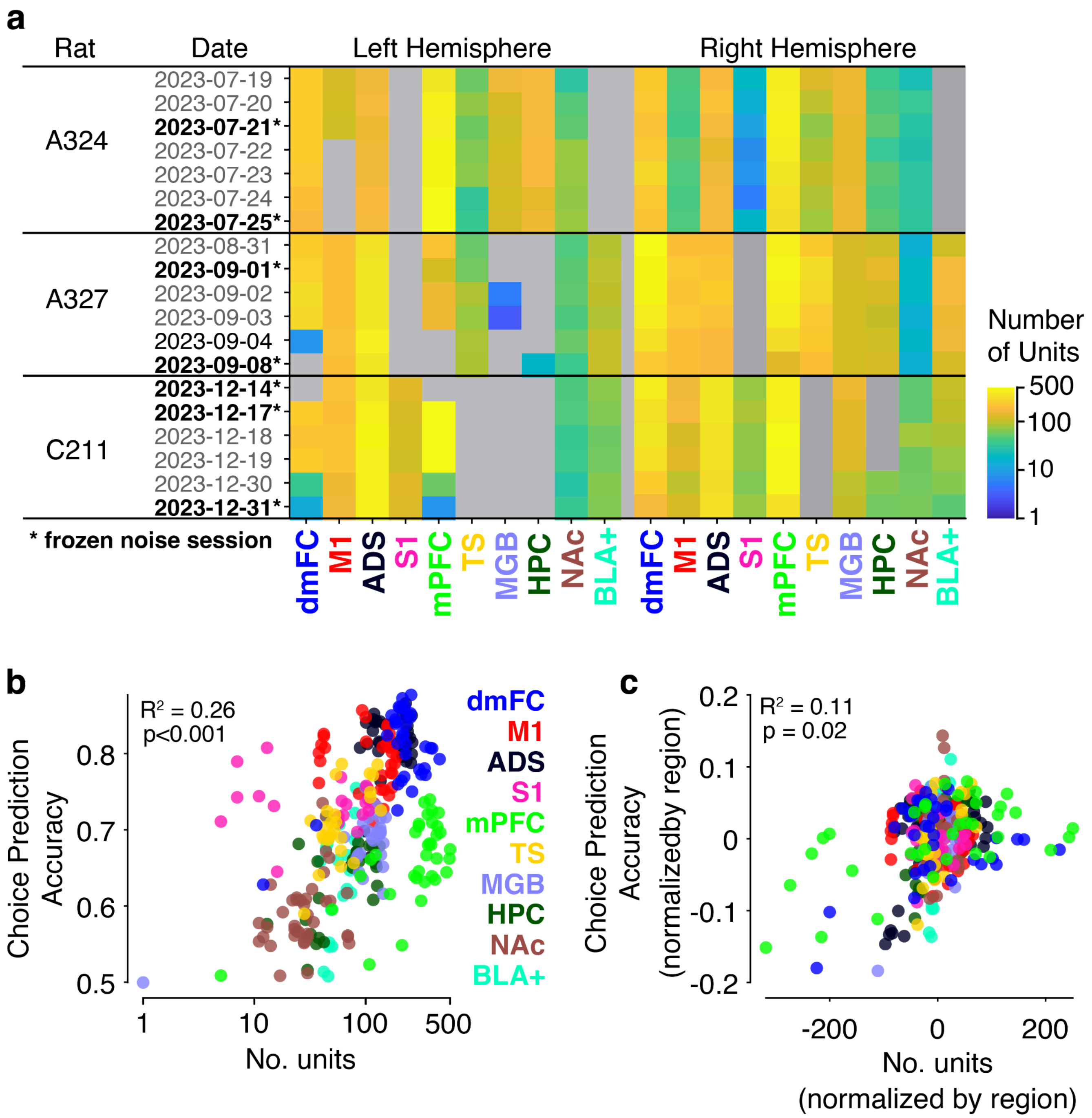


**Figure S4. Dataset breakdown and population size analyses, related to Figures 2 and 3. a**. Full breakdown showing the number of units passing quality-based inclusion criteria for each region and hemisphere, for each session in the dataset. **b**. Average choice prediction accuracy, shown separately by region and session, as a function of population size. Accuracy is averaged across timepoints from 0.7s after first click to the time of movement onset. A weak, but significant, positive relationship is observed between choice prediction accuracy and population size. The fact that datapoints corresponding to the same region cluster together indicates that this effect is largely due to real differences across regions rather than trivially being an effect of population size. Note that the region with the most recorded units was mPFC, which did not have the highest choice prediction accuracy. **c**. Same as (**b**) but after partialing-out the effect of region identity by subtracting the mean x- and y- values for each region cluster. The remaining effect of population size on choice prediction accuracy is even weaker, although not insignificant.

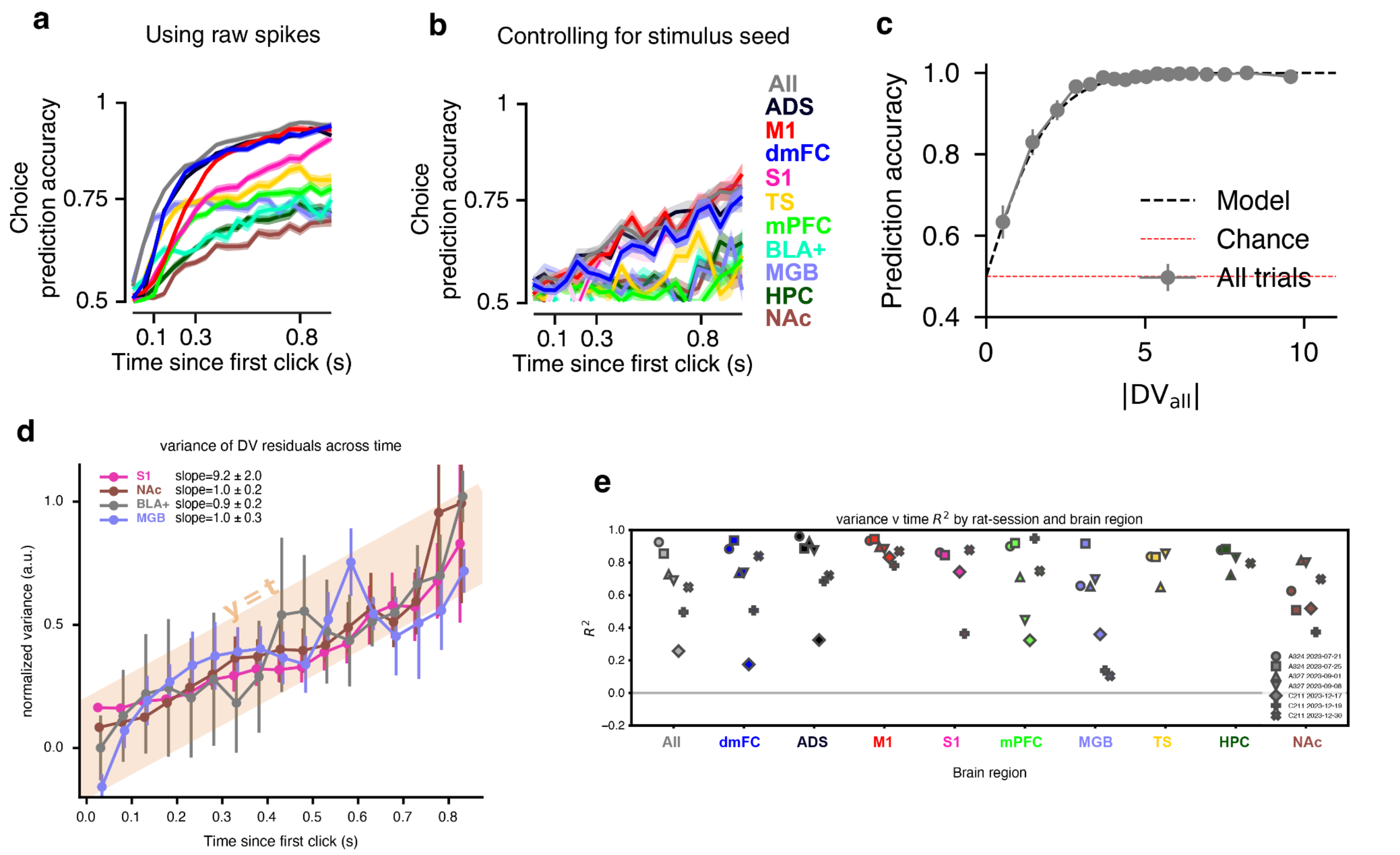


**Figure S5. DV(t) residuals reflect an ongoing evidence accumulation process, related to Figure 3. a**, Mean (+/- 1 s.e.) choice prediction accuracy as a function of time since the first click of the auditory stimulus for each of the analyzed brain regions (similar to **Figure 3d** but collapsing across hemispheres). This analysis is based on the decision variables (DV(t)s) computed from the raw spikes (i.e. not controlling for a varying stimulus across trials). 7 frozen noise sessions shown. **b**, Same as in (a) except with the mean firing rate trajectory of each neuron conditioned on stimulus (seed) i.e. subtracting the mean across groups of trials with the same click train, before performing the logistic regression. Above-chance choice decoding is still observed, indicating that fluctuations in the DV(t) independent of the stimulus are predictive of the decision. **c,** DV(t) quantitatively predicts the animal's left/right choices. Predictions from $DV_{all}$ computed from all neurons, not a single region, are shown here; results from individual regions are similar. For trials with a detected $nTc_{all}$ , here we used $DV_{all}$ at that timepoint; for trials without a detected $nTc_{all}$ , we used $DV_{all}$ at the end of the stimulus. **d,** Across-trial variance of DV(t) residuals increases linearly over time. Error bars indicate +/-1 s.e. across frozen noise seeds. Only timepoints up to 200 ms before each trial's neurally estimated time of commitment (nTc) were used.To focus on comparing linearity across regions, per-region data was shifted and scaled vertically so linear fits to each region all had an intercept of 0 and a slope of 1. Actual slopes for each region are as indicated, in units of log-odds squared per second. The regions shown here slightly but systematically deviated from a straight line and were not shown in main text FIgure 3e. **d.** Quantification of the linearity of the increase in across-trial variance of DV(t) residuals, as variance explained (R2) of a straight line fit to each session's data. Each point corresponds to one of the 7 frozen noise sessions.

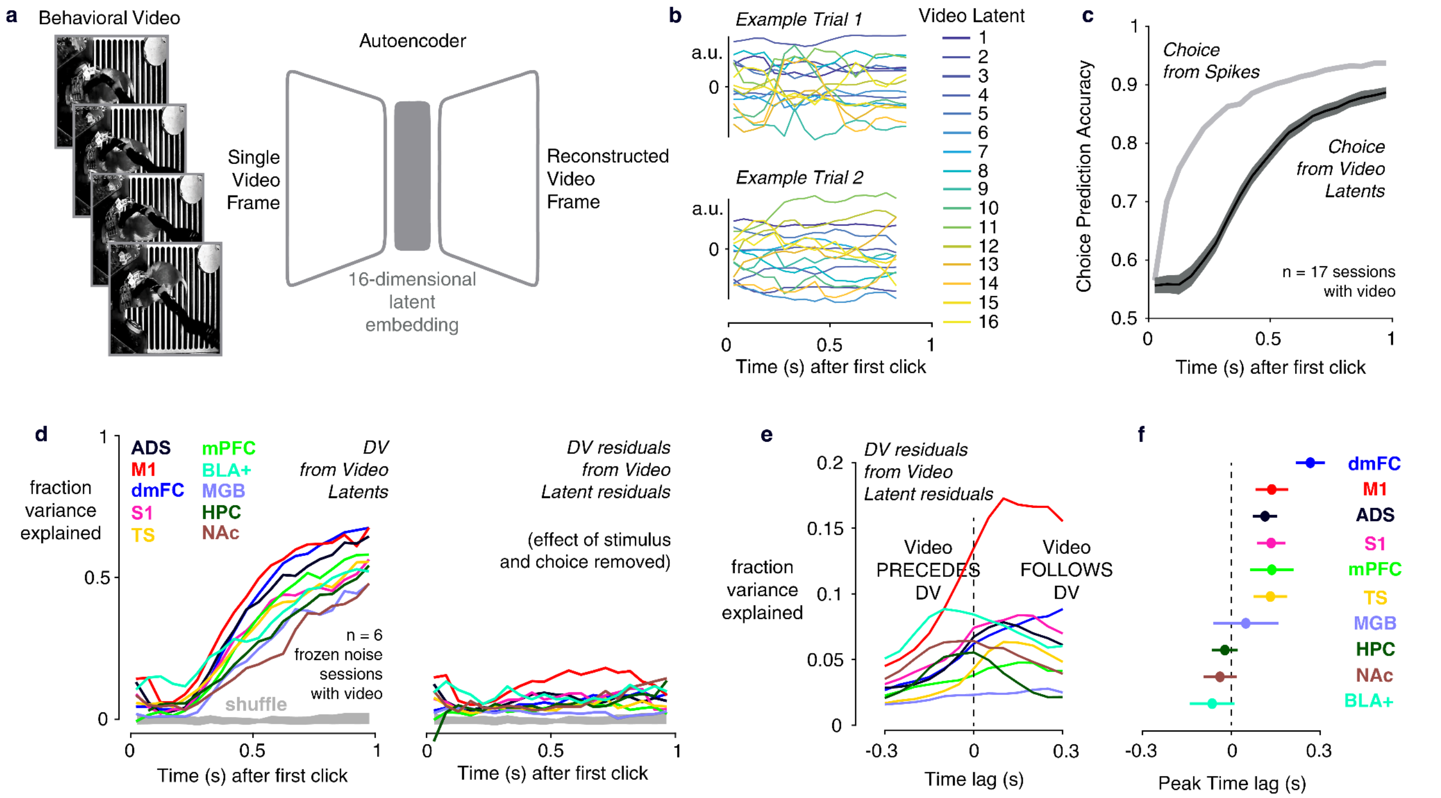


**Figure S6. DV(t) residuals and their inter-regional correlations are not driven by spontaneous movements, related to Figure 3. a.** We used a video autoencoder to obtain a 16-dimensional latent embedding of overhead behavioral video recordings that were performed in a subset of 17 behavioral sessions (see **Methods**). **b**. Example single-trial traces of the 16 video latents. **c**. Comparison of choice prediction accuracy as a function of time using either neural data (including all recorded regions) or video. Neural data contains earlier decision signals than video latents. **d**. Linear regression was used to predict DV(t)s using the 16 video latents for 6 frozen noise sessions with video recordings. Fraction of variance explained started near-zero and ramped over the course of the trial. However, when the mean within groups of trials with the same stimulus seed and choice was subtracted to obtain DV(t) and video latent residuals, the fraction of variance explained plummeted. This indicates that video is poorly able to predict DV(t)s once stimulus and choice information are removed, and therefore spontaneous movements cannot account for the DV(t) residual fluctuations. **e**. The same analysis as in the right panel of d was performed at various relative lags between the video latents and DV(t). The time-averaged fraction of variance explained is shown for each lag and for each region. **f.** The peak lag is shown for each region, with error bar indicating +/- 1 s.e. obtained from a hierarchical bootstrap across sessions. For most regions, positive lags produce the best variance explained, indicating that DV(t) residuals can be best predicted by future video for most regions. This is incompatible with DV(t) residuals being a downstream consequence of spontaneous movements.

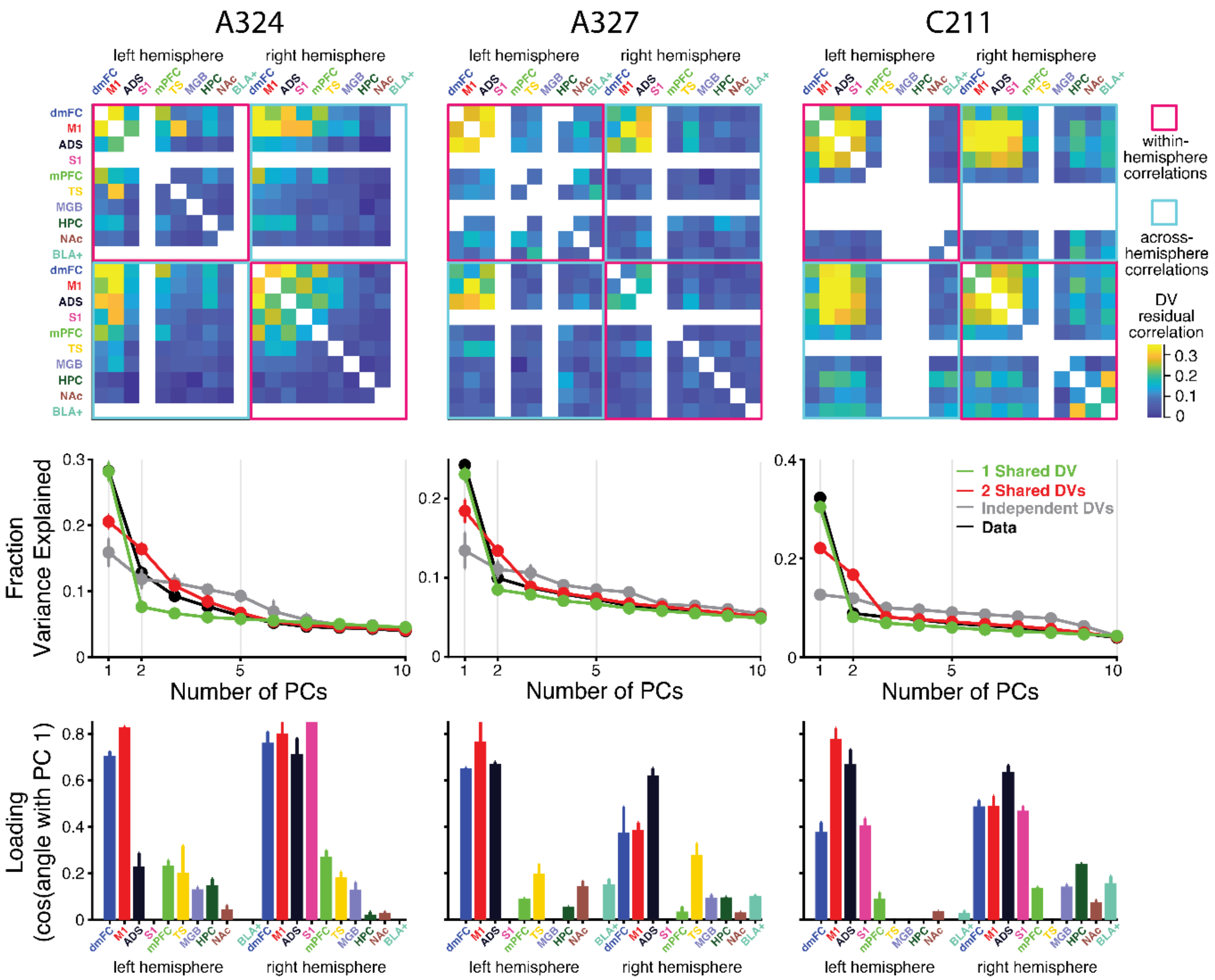


**Figure S7. DV(t) residual correlations by rat subject, related to Figure 3.** *(top)* Matrix of DV(t) residual correlations (Pearson's $\rho$) between pairs of regions, collapsed across timepoints and trials within a session. Same as **Figure 3h**, but here broken down by rat subject. *(middle)* Per-subject scree plots obtained from applying principal components analysis to the set of across-region residual DV(t)s. Each region/hemisphere pair is treated separately. The recorded data is shown along with predictions from simulations of the three competing models in **Figure 1a** (see **Methods**). Error bars indicate 1 s.e. obtained from a hierarchical bootstrap across sessions. Same as **Figure 3i** but here broken down by rat subject. *(bottom)* Plots showing the loading of each region/hemisphere pair onto the first PC. Error bars indicate 1 s.e. obtained from a hierarchical bootstrap across sessions. Same as **Figure 3j** but here broken down by rat subject.

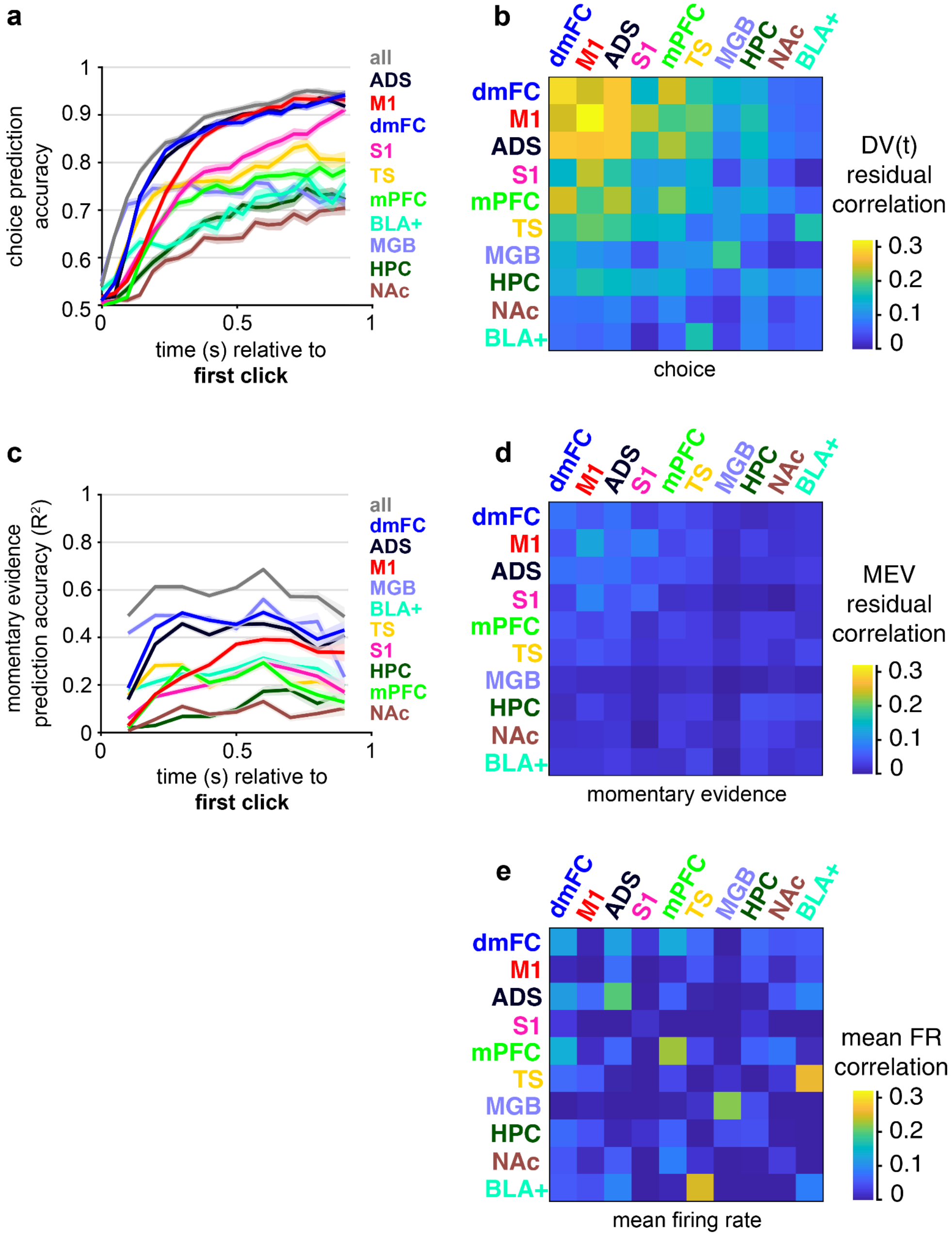


**Figure S8. Correlations between multiple scalar variables obtained from each brain region, related to Figure 3.** **a**,Time course of choice prediction accuracy using neural activity from each brain region bilaterally and from all regions combined. Mean +/- 1 s.e. across 7 frozen noise sessions shown. **b**, Matrix of DV(t) residual correlations (Pearson's $\rho$) between the between pairs of regions, across all timepoints and trials within a session. Average across the frozen noise sessions (and across the two hemispheres per session) shown. Diagonal entries correspond to the inter-hemispheric correlation for a given region. **c**, Same as (a) except shows accuracy of a linear regression model of momentary evidence using neural population activity, quantified using $R^2$. Momentary evidence is defined as the difference in right versus left clicks within a 100 ms window before the time of spiking (see **Methods**). **d**, Same as (b) except shows correlations along the dimension in neural state space best predicting momentary evidence (i.e. "the momentary evidence variable" or MEV), rather than choice. We used a shuffle-correction procedure to remove the component of correlation due to shared coding for stimulus, current choice and previous choice, as in (b). **e**, Matrix of pairwise correlations between the population mean firing rates of each pair of regions.We used a shuffle-correction procedure to remove the component of correlation due to shared coding for stimulus, current choice and previous choice, as in (b).

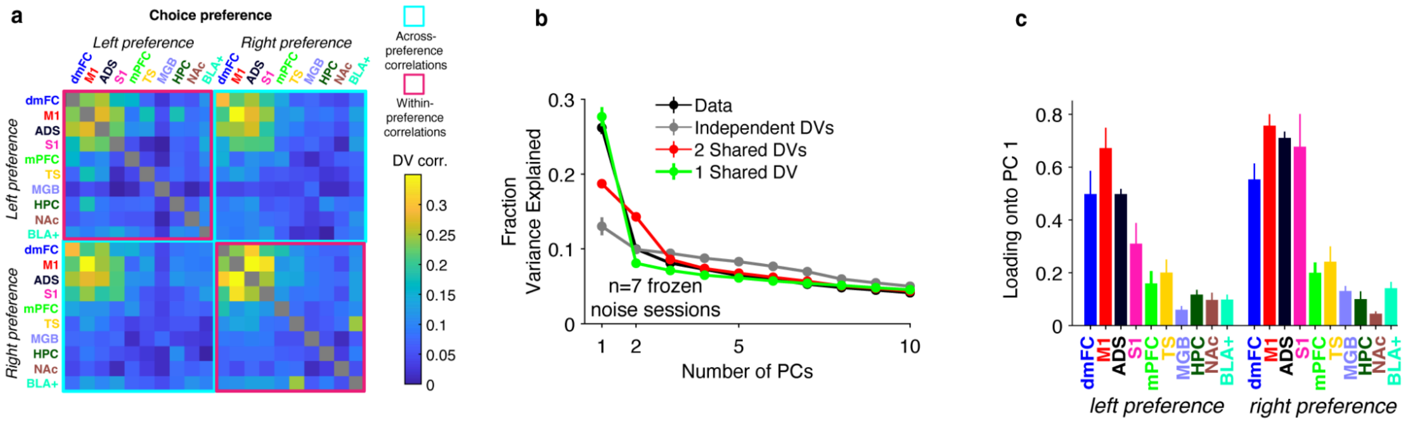


**Figure S9. DV(t) residual correlations, with regions broken down by neuronal choice preference, related to Figure 3. a)** Matrix of DV(t) residual correlations (Pearson's $\rho$) between pairs of regions, across timepoints and trials within a session. Average across the 7 frozen noise sessions shown. Same as Figure 3h, except here regions are broken down by right versus left choice preference rather than right versus left hemisphere. A similar pattern of correlations is observed. b. Scree plot obtained from applying principal components analysis to the set of residual DV(t)s. Each region/choice-preference pair is treated separately. The recorded data is shown along with predictions from simulations of the three competing models in **Figure 1a** (see **Methods**). Error bars indicate 1 s.e. obtained from a hierarchical bootstrap across sessions. The observed data is highly similar to the prediction of a single, shared DV(t), indicating left- and right-preferring subgroups of neurons across regions reflect a single shared process of evidence accumulation. c. Plots showing the loading of each region/choice-preference pair onto the first PC. Error bars indicate 1 s.e. obtained from a hierarchical bootstrap across sessions. Left- and right-preferring groups have similar loadings onto the first PC.

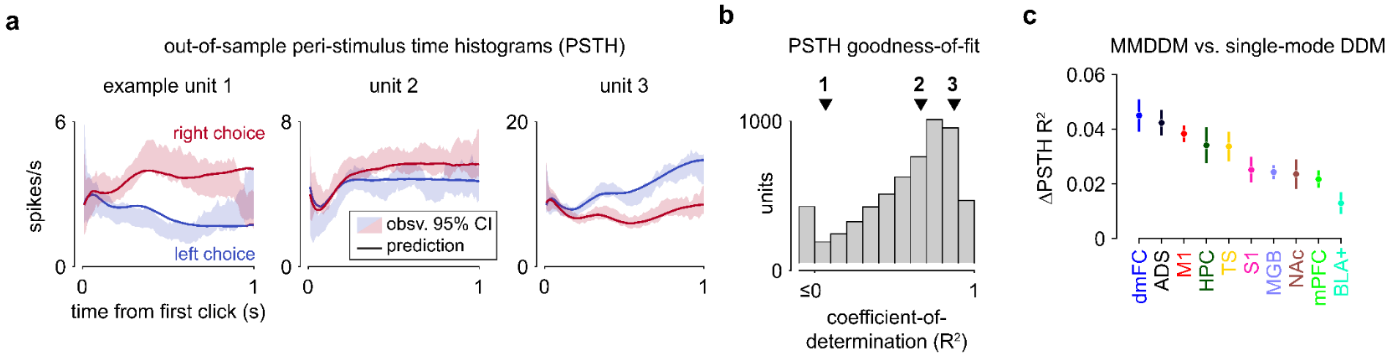


**Figure S10. MMDDM captures neuronal temporal profiles more accurately than the single-mode DDM, related to Figure 5. a**, Choice-conditioned trial-averaged responses of 3 example units. Shading indicates the 95% confidence interval of the mean across trials, and solid lines represent the mean of the cross-validated model predictions. **b,** Coefficient-of-determination ($R^2$) of the choice-conditioned PSTHs across units fit to the model, pooled across areas and sessions. **c**, Difference in the PSTH $R^2$ between MMDDM and the single-mode DDM. Data are shown as mean +/- 1 s.e. across the 19 sessions. While both models were fit to neural data pooled across regions (not separately by regions), this comparison is shown separately for each region.

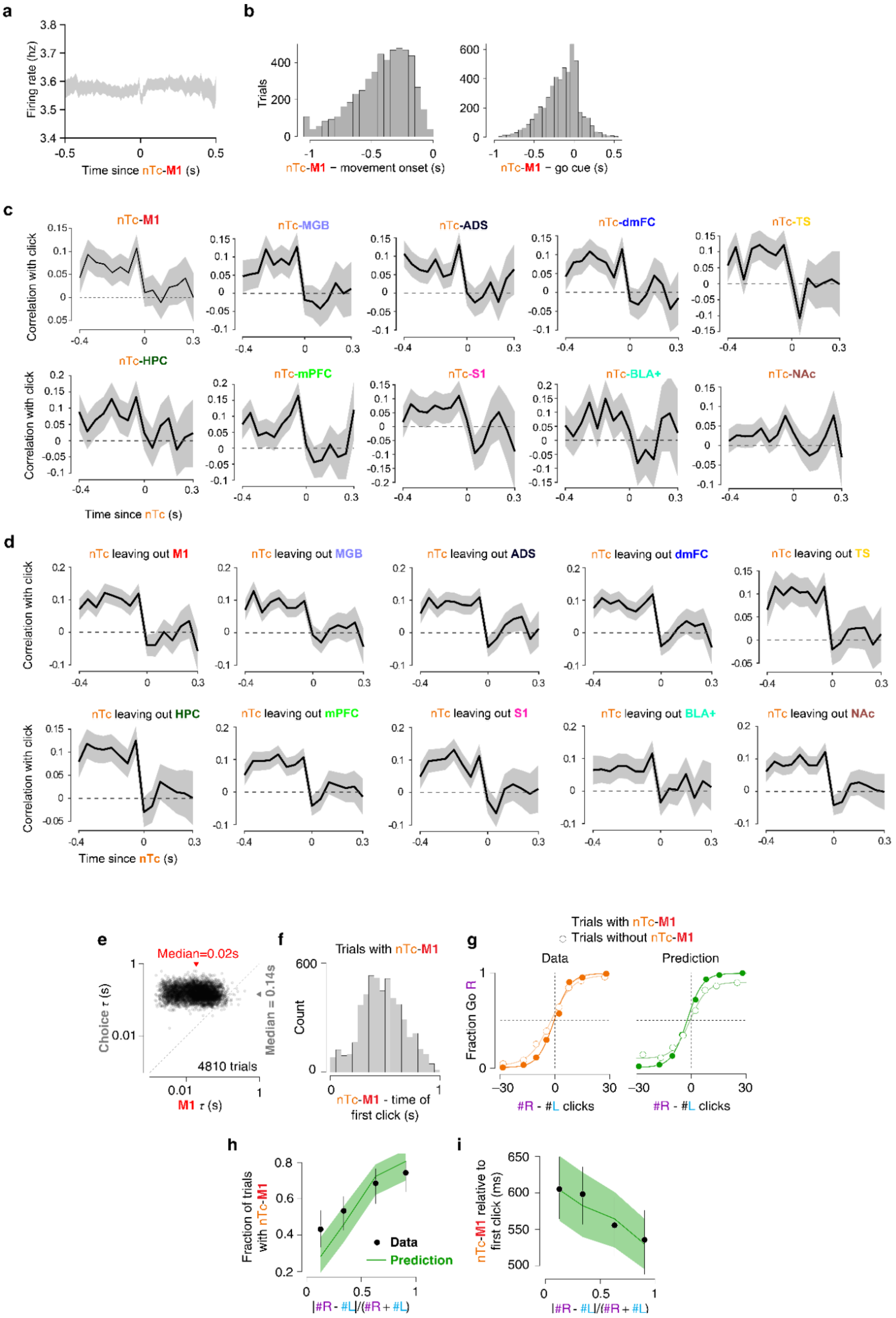
a
Firing rate (hz)
Time since nTc-M1 (s)
b
Trials
nTc-M1 − movement onset (s)
nTc-M1 − go cue (s)
c
nTc-M1
nTc-MGB
nTc-ADS
nTc-dmFC
nTc-TS
nTc-HPC
nTc-mPFC
nTc-S1
nTc-BLA+
nTc-NAc
Correlation with click
Time since nTc (s)
d
nTc leaving out M1
nTc leaving out MGB
nTc leaving out ADS
nTc leaving out dmFC
nTc leaving out TS
nTc leaving out HPC
nTc leaving out mPFC
nTc leaving out S1
nTc leaving out BLA+
nTc leaving out NAc
Correlation with click
Time since nTc (s)
e
Median=0.02s
Median = 0.14s
Choice τ (s)
4810 trials
M1 τ (s)
f
Trials with nTc-M1
Count
nTc-M1 - time of first click (s)
g
Trials with nTc-M1
Trials without nTc-M1
Data
Prediction
Fraction Go R
#R - #L clicks
h
Fraction of trials with nTc-M1
Data
Prediction
|#R - #L|/(#R + #L)
i
nTc-M1 relative to first click (ms)
|#R - #L|/(#R + #L)

**Figure S11. Characteristics and validation of nTc per region, related to Figure 5. a**, Mean firing rates across all units, averaged across trials aligned to nTc-M1. Shading indicates 95% bootstrapped confidence of the trial-averaged mean (n=5,135 trials). **b**, Left, distribution of nTc across trials that can be detected using M1 spikes, with respect to the onset of the animal's choice-reporting movement. Right, same as left, but with respect to the go cue. **c**, Weight of stimulus cues on the animal's decision choice ("psychophysical kernel"), relative to the nTc inferred using neural activity from different brain regions. For all regions except NAc, this test validates the neurally-estimated nTc by showing that cues before, but not after, nTc, affect choice. **d**, Psychophysical kernel estimated by aligning to the nTc inferred in a leave-one-region-out manner. Each plot shows the kernel aligned to an nTc estimate that excluded spikes from one region. **e,** the timing precision of the estimated nTc is much greater when spikes are used to infer nTc than compared to using the animal's choice instead. **f**, nTc-M1 is broadly distributed relative to stimulus start. Similar results are found for other brain regions (not shown). **g,** trials with nTc show higher performance compared to those without, similarly in the experimental data and in the MMDDM model used to infer nTc. **h,** Easier trials have a higher fraction of trials with a detected nTc, similarly in data and MMDDM model. **i,** Easier trials have earlier nTcs, similarly in data and MMDDM model.

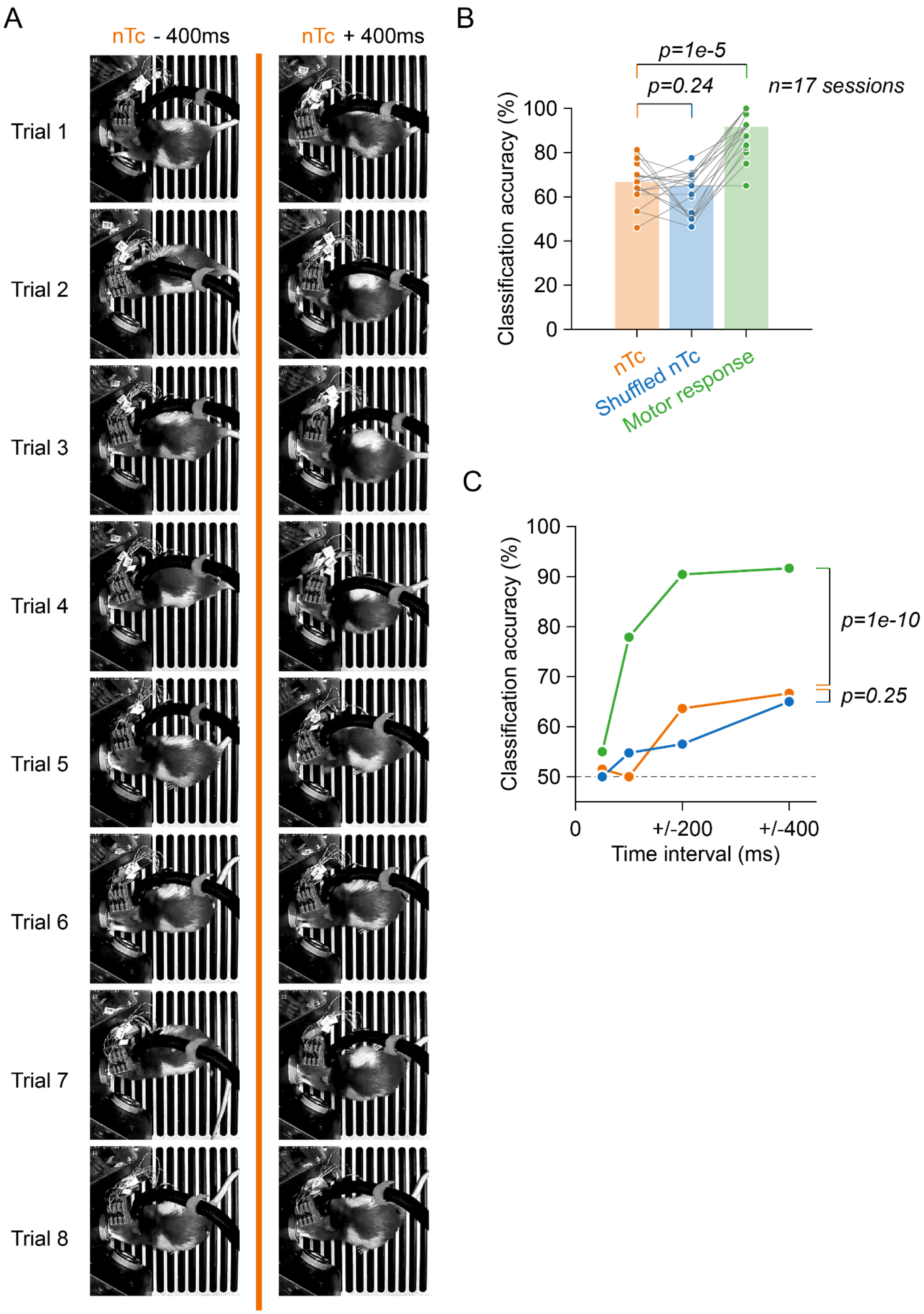


**Figure S12. Animal postures captured before and after nTc could not be distinguished by a convolutional neural network, related to Figure 5.** A) Representative video frames captured 400 ms before (left) and 400 ms after (right) the neurally-inferred time of commitment (nTc) across eight example trials. Only trials where the animal's nose-out movement response occurred >400 ms after nTc were included. B) Decoding accuracy of nTc-centered frames. A convolutional neural network (SqueezeNet) was trained to classify frames as "pre-nTc" or "post-nTc." Bar heights represent the median accuracy across n=17 sessions. "Shuffled nTc" serves as a negative control, while "Motor response" (classifying frames around the rat's exit from the center port) serves as a positive control. C) Effect of temporal window on classification.

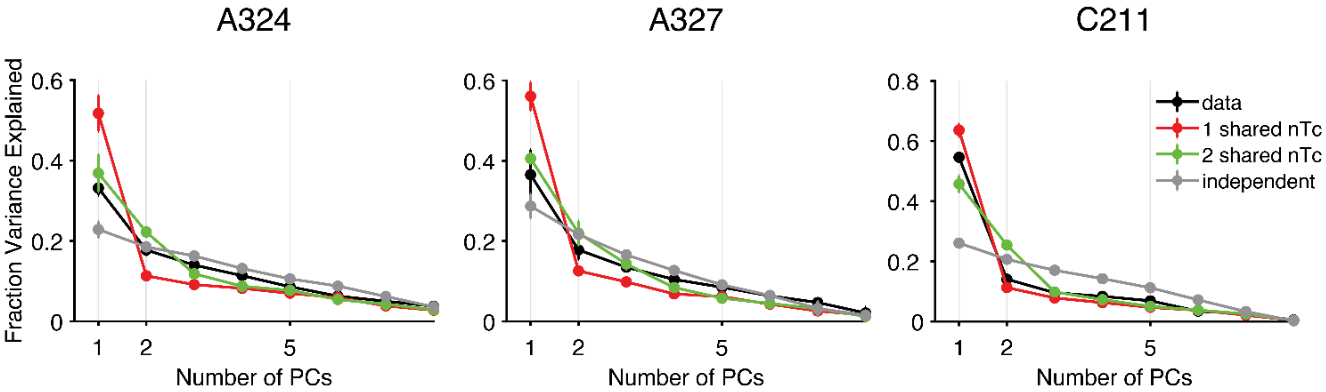


**Figure S13. nTc Residuals dimensionality broken down by rat subject, related to Figure 6.** Scree plots produced from applying probabilistic PCA to the set of across-region nTc residuals, generated from simulations implementing the competing hypotheses in Figure 6a (see **Methods**) as well as using the observed data. Error bars indicate 1 s.e. using a hierarchical bootstrap across sessions**.** Same as main text Figure 6c,d but broken down by rat subject.

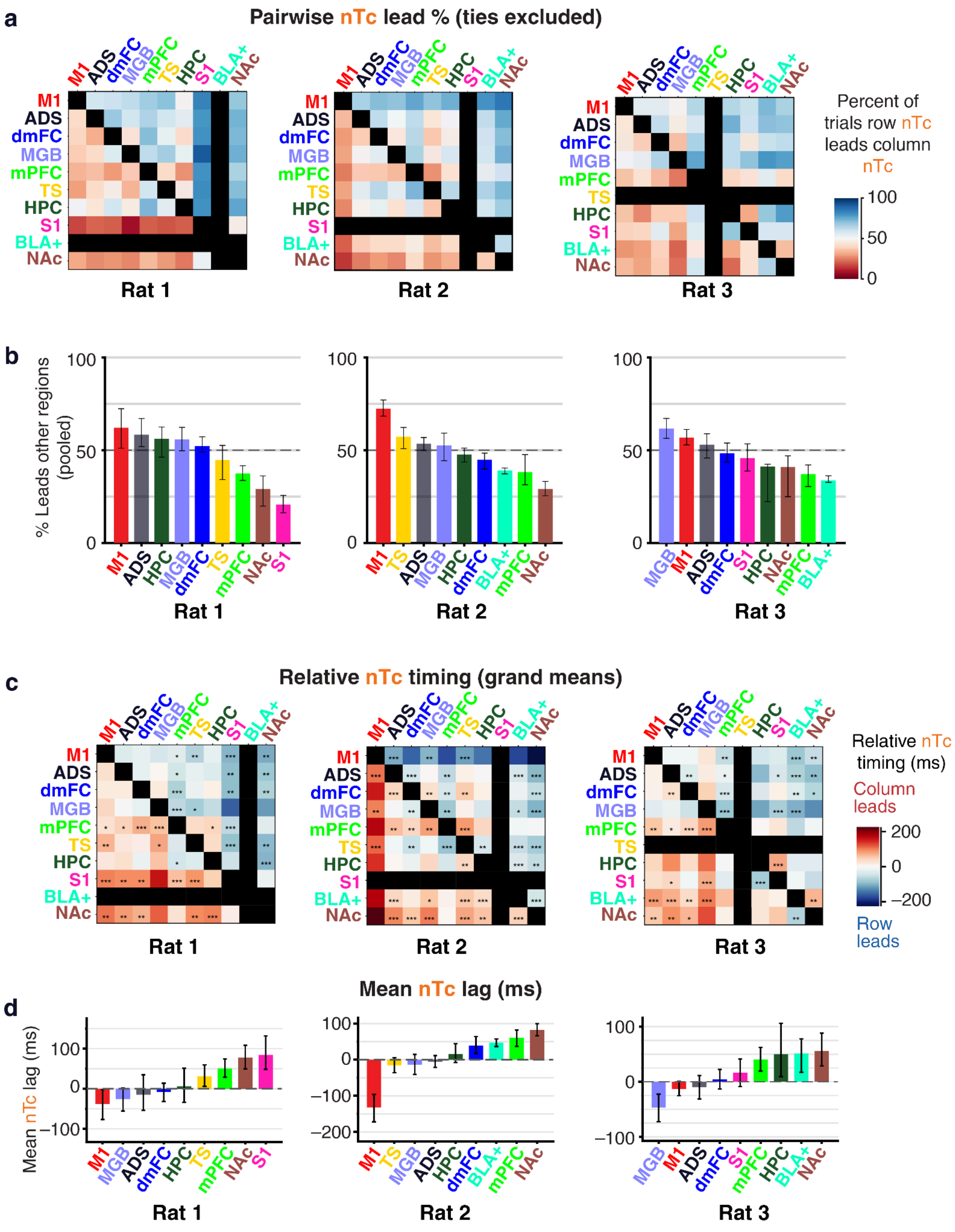


**Figure S14. Relative nTc timing broken down by rat subject, related to Figure 6. a**, Matrices showing the proportion of trials for which the row and column regions both had an nTc in which the row region's nTc happened first. Trials in which the pair had tied nTc times were excluded from analysis. Proportions were calculated on trials pooled across sessions within each rat. **b**, Mean proportion of trials in which each region led others, calculated as the percentage pooled over rows of the matrix shown in (a). Error bars are the bootstrapped 95% confidence interval. **c**, Matrices representing the trial-averaged difference in nTc timing between all pairs of brain regions, for each rat. Choice AUC distributions were matched before calculating nTc (see Methods). Each pairwise comparison includes only trials with nTc detected in both regions. Negative values indicate the row region's nTc tends to lead the column region's nTc. Statistics were performed using a linear mixed model with a random intercept for recording session (***, $p < 0.001$; **, $p < 0.01$; *, $p < 0.05$). Black squares indicate missing data. Bottom row, Mean of nTc lags for each brain region, weighted within session by the number of trials in each pairwise comparison. Error bars are the bootstrapped 95% CIs over sessions. **d**, Each region's mean nTc lag relative to all others, corresponding to the mean of each row in the matrix in panel c.

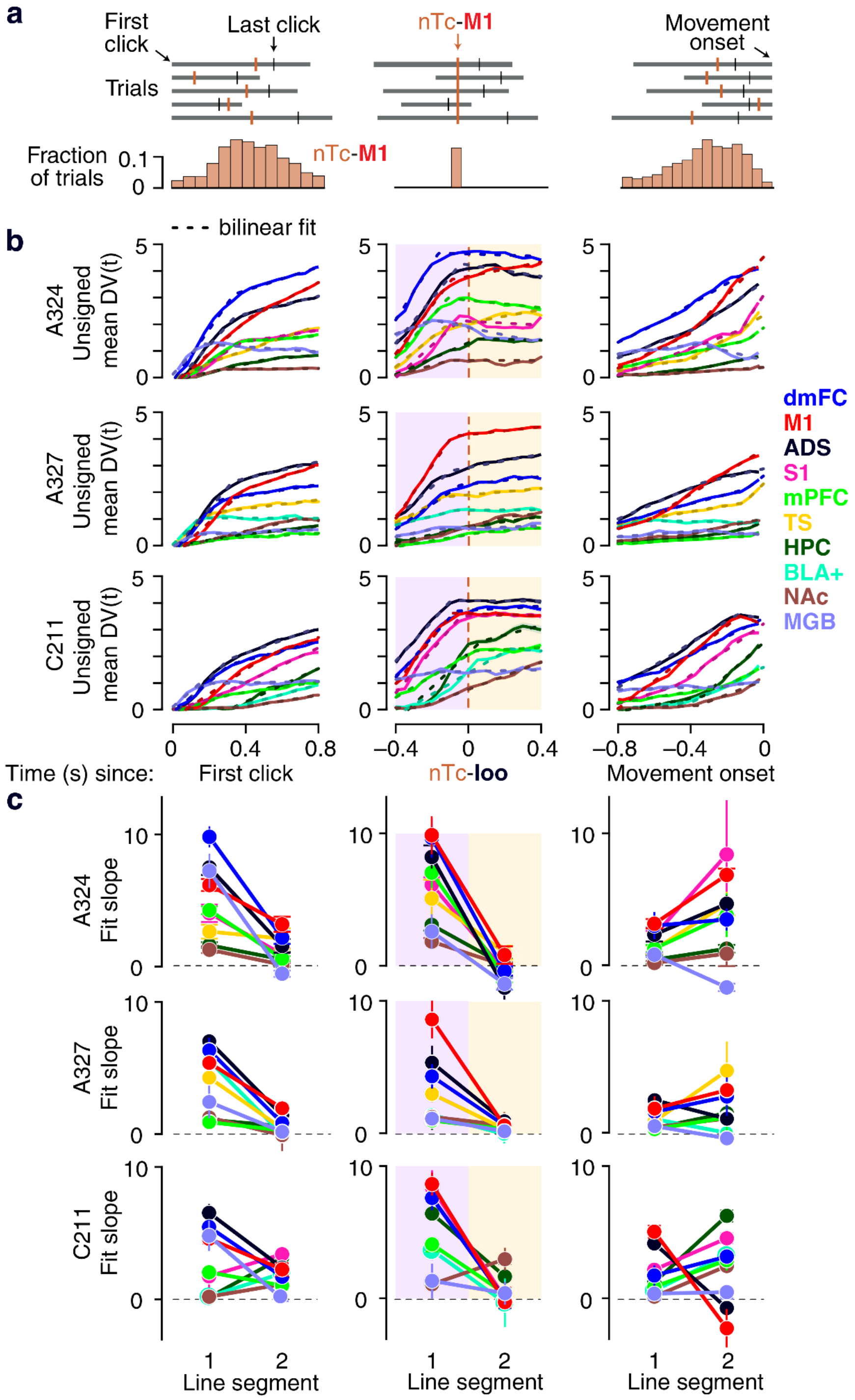


**Figure S15. Bilinear fits to DV(t) traces for various task event alignments, broken down by rat subject, related to Figure 7.** **a**, Timing of nTc-M1 relative to key trial events. Top, sequence of the first click, nTc-M1, and movement onset for 5 example trials. Bottom, distribution of nTc-M1 for each of these three events across all trials. Only trials with valid estimates of nTc-M1 are included. **b**, Trial-averaged DV(t) traces from multiple brain regions, with sign flipped for left choice trials, and aligned to each of three trial events. For the nTc-aligned data, as in the main text, each region's data was aligned to an estimate of nTc excluding spikes from that region, to avoid circularity. Each row corresponds to one of the three rats. **c**, Slopes of the first and second line segments of the bilinear fits to the traces shown in b. Each row corresponds to one of three rats.

| *ion name and abbreviation (Paxino Watson Rat Brain Atlas)* | *n name and abbreviation in this* | *corded in Ra* | *corded in Ra* | *corded in Ra* |
|---|---|---|---|---|
| caudate putamen/striatum (CPu) | anterior dorsal striatum (ADS) | L,R | L, R | L, R |
| caudate putamen/striatum (CPu) | tail of the striatum (TS) | L, R | L, R | L, R |
| primary motor cortex (M1) | primary motor cortex (M1) | L, R | L, R | L, R |
| cingulate cortex, area 1 (Cg1) | dorsomedial frontal cortex (dmFC) | L, R | L, R | L, R |
| secondary motor cortex (M2) | dorsomedial frontal cortex (dmFC) | L, R | L, R | L, R |
| prelimbic cortex (PrL) | medial prefrontal cortex (mPFC) | L, R | L, R | L, R |
| medial orbital cortex (MO) | medial prefrontal cortex (mPFC) | X | L, R | L |
| primary somatosensory cortex (S1) | primary somatosensory cortex (S1) | L, R | L, R | L, R |
| dentate gyrus (DG) | hippocampus (HPC) | L, R | L, R | R |
| field CA1 of the hippocampus (CA1) | hippocampus (HPC) | R | L, R | R |
| accumbens nucleus (Acb) | nucleus accumbens (NAc) | L, R | L, R | L, R |
| subbrachial nucleus (SubB) | subbrachial nucleus (SBN) | L, R | R | R |
| medial geniculate nucleus (MG) | medial geniculate body (MGB) | L, R | L, R | R |
| basolateral amygdaloid nucleus (BL) | basolateral amygdala+ (BLA+) | X | L, R | X |
| Dorsal endopiriform nucleus (DEn) | basolateral amygdala+ (BLA+) | X | X | L, R |
| peripeduncular nucleus (PP) | peripeduncular nucleus (PP) | X | L | X |
| piriform cortex (Pir) | piriform cortex (Pir) | X | L, R | X |
| globus pallidus (GP) | globus pallidus (GP) | L | X | X |
| primary visual cortex (V1) | primary visual cortex (V1) | L, R | L, R | R |
| substantia nigra, reticular part (SNR) | substantia nigra (SN) | X | L | X |
| dorsal subiculum (DS) | dorsal subiculum (DS) | L | X | X |

**Supplementary Table 1. Complete listing of recorded brain regions.** In some cases, we refer to a brain region differently than what is used in the Paxinos and Watson atlas (Paxinos, G. & Watson, C. The Rat Brain in Stereotaxic Coordinates. Elsevier, 2009). In other cases, we combine two regions named by Paxinos and Watson and refer to both by one name. ‘L’ indicates that the region was recorded from the left hemisphere of a rat subject; ‘R’ indicates it was recorded from the right hemisphere. Cells of the table marked ‘X’ were not recorded from in the given subject. Units appearing to be in BL and DEn were grouped into ‘BLA+’ to indicate our uncertainty in the true identity of units recorded in these adjacent areas.
